# Probabilistic Harmonization and Annotation of Single-cell Transcriptomics Data with Deep Generative Models

**DOI:** 10.1101/532895

**Authors:** Chenling Xu, Romain Lopez, Edouard Mehlman, Jeffrey Regier, Michael I. Jordan, Nir Yosef

**Author notes:** These authors contributed equally to this work (Chenling Xu, Romain Lopez, Edouard Mehlman). Corresponding author. (N.Y.).

## Abstract

As single-cell transcriptomics becomes a mainstream technology, the natural next step is to integrate the accumulating data in order to achieve a common ontology of cell types and states. However, owing to various nuisance factors of variation, it is not straightforward how to compare gene expression levels across data sets and how to automatically assign cell type labels in a new data set based on existing annotations. In this manuscript, we demonstrate that our previously developed method, scVI, provides an effective and fully probabilistic approach for joint representation and analysis of cohorts of single-cell RNA-seq data sets, while accounting for uncertainty caused by biological and measurement noise. We also introduce single-cell ANnotation using Variational Inference (scANVI), a semi-supervised variant of scVI designed to leverage any available cell state annotations — for instance when only one data set in a cohort is annotated, or when only a few cells in a single data set can be labeled using marker genes. We demonstrate that scVI and scANVI compare favorably to the existing methods for data integration and cell state annotation in terms of accuracy, scalability, and adaptability to challenging settings such as a hierarchical structure of cell state labels. We further show that different from existing methods, scVI and scANVI represent the integrated datasets with a single generative model that can be directly used for any probabilistic decision making task, using differential expression as our case study. scVI and scANVI are available as open source software and can be readily used to facilitate cell state annotation and help ensure consistency and reproducibility across studies.

## Introduction

Recent technological improvements in microfluidics and low volume sample handling [1] have enabled the emergence of single-cell transcriptomics [2, 3] as a popular tool for analyzing biological systems [4, 5, 5]. This growing popularity along with a continued increase in the scale of the respective assays [7] has resulted in massive amounts of publicly available data and motivated large scale community efforts such as the Human Cell Atlas [8], Tabula Muris [9] and the BRAIN Initiative Cell Census Network [10]. The next natural step in the evolution of this field is therefore to integrate many available datasets from related tissues or disease models in order to increase statistical robustness [11], achieve consistency and reproducibility among studies [12, 13], and ultimately converge to a common ontology of cell states and types [8, 14].

A fundamental step toward the ideal of a common ontology is data *harmonization*, namely integration of two or more transcriptomics datasets into a single dataset on which any downstream analysis can be applied. We use the term harmonization rather than *batch effect correction* in order to emphasize that the input datasets may come from very different sources (*e.g.*, technology, laboratory), and from samples with a different composition of cell types. A wide range of methods have already been developed for this fundamental problem, initially for Microarrays and later on for bulk RNA sequencing, such as ComBat [15] and limma [16] which rely on generalized linear models with empirical Bayes shrinkage to avoid over-correction. More recently, similar methods have been proposed specifically for single-cell RNA sequencing (scRNA-seq), such as ZINB-WaVE [17], which explicitly accounts for the overabundance of zero entries in the data. However, because of their linear assumptions, these approaches may not be appropriate when provided with a heterogeneous sample that includes different cell states, each of which may be associated with a different sample-to-sample bias [12]. With these limitations in mind, the next generation of methods turned to non-linear strategies. Broadly speaking, each of these methods includes a combination of two components: (i) joint factorization of the input matrices (each corresponding to a different dataset) to learn a joint low-dimensional latent representation. This is usually done with well established numerical methods, such as integrative non-negative matrix factorization (LIGER [18]), singular value decomposition (Scanorama [19]), or canonical correlation analysis (Seurat Alignment [13]); (ii) additional non-linear transformation of the resulting latent representations so as to optimally “align” them onto each other. This is usually done using heuristics, such as alignment of mutual nearest neighbors (MNN [12], Scanorama [19] and Seurat Anchors [20]), dynamic time warping (Seurat Alignment [13]) or quantile normalization (LIGER [18]). While this family of methods has been shown to effectively overlay different datasets, it suffers from two important limitations. First, an explicit alignment procedure may be difficult to tune in a principled manner and consequently result in over-normalization, especially in challenging cases where the cell type composition is different between datasets and when technical differences between samples are confounded with biological differences of interest. Second, the alignment is done in an ad hoc manner and lacks probabilistic interpretability. Consequently, the resulting harmonized dataset is of limited use and cannot be directly applied for differential expression and other probabilistic decision-making tasks.

Another recent line of work makes use of neural networks to learn a joint representation of multiple datasets (SAUCIE [21]) or project one dataset into another (maximum mean discrepancy [MMD] ResNet [22]). These methods rely on an explicit non-parametric measure of discrepancy between probability distributions (MMD) to match either the latent spaces or directly the gene expression values from pairs of datasets. However, using the MMD with a universal kernel explicitly assumes that the cell type proportion is similar in all the datasets, which may be less suitable in the general case of data harmonization.

Besides harmonization, another important and highly related problem is that of automated *annotation* of cell state. In principle, there are two ways to approach this problem. The first is *ab initio* labeling of cells based on marker genes or gene signatures [13, 23, 23]. While this approach is intuitive and straightforward, its performance may be affected in the plausible case where marker genes are absent due to limitations in sensitivity. The second approach is to “transfer” annotations between datasets. In the simplest scenario, we have access to one dataset where states have been annotated either *ab initio*, or using additional experimental measurements (e.g., protein expression [3, 25] or lineage tracing [26]) and another, unannotated dataset from a similar condition or tissue. The goal is to use the labeled data to derive similar annotations for the second dataset, whenever applicable. This task is often complicated by factors such as differences in technology (e.g., using Smart-Seq2 data to annotate 10x Chromium data), partial overlap in cell type composition (i.e., not all labels should be transferred and not all unannotated cells should be assigned a label), complex organization of the labels (e.g., hierarchy of cell types and sub-types [27], continuum along phenotypic or temporal gradients), partial labeling (i.e., only a subset of cells from the “annotated” dataset can be assigned a label confidently), and the need to handle multiple (more than 2) datasets in a principled and scalable manner. One way to address the annotation problem with this approach is learning a classifier [27, 28] in order to predict a fixed stratification of cells. However, this approach might be sensitive to batch effects, which could render a classifier based on a reference dataset less generalizable to an unannotated dataset. Another, more flexible approach is to transfer annotations by first harmonizing the annotated and unannotated datasets, thus also gaining from the benefits of having a single dataset that can be subject to additional, joint, downstream analysis.

In this paper, we propose a strategy to address several of the outstanding hurdles in both of the harmonization and annotation problems. We first demonstrate that single-cell Variational Inference (scVI) [29] a deep generative model we previously developed for probabilistic representation of scRNA-seq data — performs well in both harmonization and harmonization-based annotation, going beyond its previously demonstrated capacity to correct batch effects. We then introduce single-cell ANnotation using Variational Inference (scANVI), a new method that extends scVI and provides a principled way to address the annotation problem probabilistically while leveraging any available label information. scANVI uses a semi-supervised generative model, which can be utilized for both approaches to the annotation problem. In the first scenario, we are concerned with a single dataset in which only a subset of cells can be confidently labeled (e.g., based on expression of marker genes) and annotations should then be transferred to other cells, when applicable. In the second scenario, annotated datasets are harmonized with unannotated datasets and then used to assign labels to the unannotated cells.

The inference procedure for both of the scVI and scANVI models relies on neural networks, stochastic optimization and variational inference [30, 31] and scales to large numbers of cells and datasets. Furthermore, both methods provide a complete probabilistic representation of the data, which non-linearly controls not only for sample-to-sample bias but also for other technical factors of variation such as over-dispersion, library size discrepancies and zero-inflation. As such, each method provides a single probabilistic model that underlies the harmonized gene expression values (and the cell annotations, for scANVI), and can be used for any type of downstream hypotheses testing. We demonstrate the latter point through a differential expression analysis on harmonized data. Furthermore, through a comprehensive analysis of performance in various aspects of the harmonization and annotation problems and in various scenarios, we demonstrate that scVI and scANVI compare favorably to current state-of-the-art methods.

## Results

In the following we demonstrate that our framework compares favorably to the existing state of the art in the harmonization and annotation problems in terms of accuracy, scalability, and adaptability to various settings. The first part of the paper focuses on the harmonization problem and covers a range of scenarios, including harmonization of datasets with varying levels of biological overlap, handling cases where the data is governed by a continuous (e.g., pseudotime) rather than discrete (cell types) form of variation, and processing multiple (> 20) datasets. While we demonstrate that scVI performs well in these scenarios, we also demonstrate that the latent space leaned by scANVI provides a proper harmonized representation of the input datasets — a property necessary for guaranteeing its performance in the annotation problem. In the second part of this manuscript we turn to the annotation problem and study its two main settings, namely transferring labels between datasets and *ab-inito* labeling. In the first setting we consider the cases of datasets with a complete or partial biological overlap and use both experimentally- and computationally-derived labels to evaluate our performance. In the second settings, we demonstrate how scANVI can be used effectively to annotate a single dataset by propagating high confidence seed labels (based on marker genes) and by leveraging a hierarchical structure of cell state annotations. Finally, we demonstrate that the generative models inferred by scANVI and scVI can be directly applied for hypotheses testing, using differential expression as a case study.

### Joint modeling of scRNA-seq datasets

We consider *K* different scRNA-seq datasets (Figure 1a). A dataset indexed by *k* ∈ {1, …, *K*} consists of a *N*_*k*_ × *G*_*k*_ matrix where each entry records the number of transcripts observed for each of *G*_*k*_ genes in each of *N*_*k*_ cells. In this work, we use a standard heuristic to filter the genes and generate a common (possibly large) gene set of size *G* (**Online Methods**), and thus format our data as a unique (Σ_*k*_ *N*_*k*_) × *G* matrix whose individual entries are noted *x*_*ng*_ — namely the expression of gene *g* in cell *n*. We use *s*_*n*_ ∈ {1, …, *K*} to represent the dataset from which each cell *n* was generated. Furthermore, a subset of the cells may be associated with a cell state annotation *c*_*n*_ ∈ *C*, which can describe either discrete cell types or hierarchical cell types. More complex structures over labels such as gradients are left as a future research direction.

**Figure 1:**
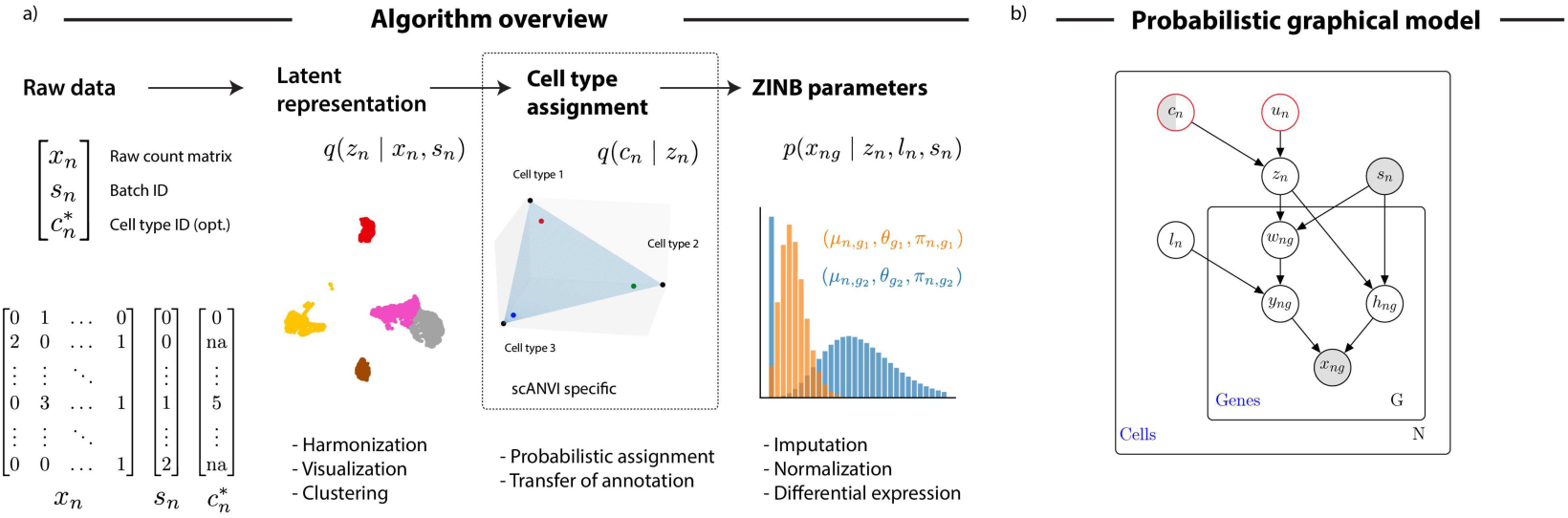
Harmonization of scRNA-seq datasets with generative models. (*a*) Schematic diagram of the variational inference procedure in both of the scVI and scANVI models. We show the order in which random variables in the generative model are sampled and how these variables can be used to derive biological insights. (*b*) The graphical models of scVI and scANVI. Vertices with black edges represent variables in both scVI and scANVI, and vertices with red edges are unique to scANVI. Shaded vertices represent observed random variables. Semi-shaded vertices represent variables that can be either observed or random. Empty vertices represent latent random variables. Edges signify conditional dependency. Rectangles (“plates”) represent independent replication. The complete model specification and definition of internal variables is provided in the **Online Methods**.

Since the problem of data harmonization of single-cell transcriptomics is difficult and can potentially lead to over-correction (Supplementary Figure 1) [32], we propose a fully-generative method as a robust and principled approach to address it. In our previous work [29], we built single-cell Variational Inference (scVI), a deep generative model where the expression level *x*_*ng*_ is zero-inflated negative binomial (ZINB) when conditioned on the dataset identifier (*s*_*n*_), and two additional latent random variables. The first, which we denote by *l*_*n*_, is a one-dimensional Gaussian accounting for the variation in capture efficiency and sequencing depth. The second is *z*_*n*_ is a low dimensional Gaussian vector that represents the remaining variability (Figure 1b). This vector is expected to reflect biological differences between cells, and can be effectively used for visualization, clustering, pseudotime inference and other tasks. Since the scVI model explicitly conditions on the dataset identifier, it provides an effective way of controlling for technical sample-to-sample variability. However, scVI is unsupervised and does not make use of the available annotations *c*_*n*_, which can further guide the inference of an informative latent representation *z*_*n*_. To this end, we present a more refined hierarchical structure for *z*_*n*_. We draw *z*_*n*_ as a mixture conditioned on the cell annotation *c*_*n*_ and another latent variable *u*_*n*_, accounting for further biological variability within a cell type (**Online Methods**). We name the resulting approach single-cell ANnotation using Variational Inference (scANVI). Notably, our work has substantial overlap with machine learning research such as semi-supervised learning in the context of variational-autoencoders [33, 34] domain adaptation [35] and label propagation [36]. We review these related research topics in **Supplementary Note 1**.

The variables *z*_*n*_, inferred either with scVI or scANVI, provide an embedding of all cells in a single, joint latent space. Since this latent space is inferred while controlling for the dataset of origin (*s*_*n*_), it inherently provides a way to address the harmonization problem. The annotation of unlabeled cells can therefore be conducted with scVI using their proximity to annotated cells in the joint latent space (e.g., using majority vote over the *k*-nearest neighbors). The scANVI model provides a more principled way to annotate cells, namely through a Bayesian semi-supervised approach. Once fitted, the model is able to provide posterior estimates for the unobserved cell state *c*_*n*_, which can be particularly useful when labels cannot be entirely trusted. Because the marginal distribution *p*(*x*_*ng*_, *c*_*n*_ | *s*_*n*_) if *c*_*n*_ observed (resp. *p*(*x*_*ng*_ | *s*_*n*_) otherwise) is not amenable to exact Bayesian computation, we use variational inference parameterized by neural networks to approximate it [30] (**Online Methods**).

Notably, scANVI and scVI both have a certain number of hyperparameters. In the following evaluations, conducted on different datasets and different scenarios, we use the exact same set of hyperparameters in order to demonstrate that our methods can be applied with a minimal requirement of hyperparameter tuning (**Online Methods**). We provide a robustness study for hyperparameters in the context of harmonization in Supplementary Figure 2. We further discuss the underlying assumptions of our framework in the context of competing harmonization methods in **Supplementary Note 2**.

### Datasets

We apply our method on datasets generated by a range of technologies (10x Chromium [3, 37], plate-based Smart-Seq2 [38], Fluidigm C1 [39], MARS-Seq [40], inDrop [41] and CEL-Seq2 [42]), spanning different numbers of cells (from a few thousand to over a hundred thousand cells), and originating from various tissues (mouse bone marrow, human peripheral mononuclear blood cells [PBMCs], human pancreas, mouse brain). Datasets are listed and referenced in Supplementary Table 1.

### Harmonizing pairs of datasets with a discrete population structure

We conducted a comparative study of harmonization algorithms on four different instances, each consisting of a pair of datasets. The first pair (PBMC-CITE [25], PBMC-8K [37]) represents the simplest case, in which the two datasets come from very similar biological settings (i.e., PBMCs) and are generated by the same technology (i.e., 10x) but in different labs (i.e., akin to batch correction). A second scenario is that of similar tissue but different technologies, which we expect to be more challenging as each technology comes with its own characteristics and biases [43]. For instance, some methods (10x, CEL-Seq2) profile the end of the transcript and use Unique Molecular Identifier (UMI) to mitigate inflation in counting, whereas others (e.g., most applications of Smart-Seq2) consider the full length of the transcript without controlling for this potential bias. Additionally, some protocols (e.g., Smart-Seq2) tend to have higher sensitivity and capture more genes per cell compared to others. Finally, studies using droplet based protocols tend to produce much larger numbers of cells compared to plate-based methods. We explore three such cases, including a bone marrow 10x and Smart-Seq2 pair from the Tabula Muris project (MarrowTM-10x, MarrowTM-ss2 [9]), a pancreas inDrop and CEL-Seq2 pair (Pancreas-InDrop, Pancreas-CEL-Seq2 [44]), and a dentate gyrus 10x and Fluidigm C1 pair (DentateGyrus-10x, DentateGyrus-C1 [45]).

Successful harmonization should satisfy two somewhat opposing criteria (Supplementary Figure 1). On the one hand, cells from the different datasets should be well mixed; namely, the set of *k*-nearest neighbors (*k*NN) around any given cell (computed e.g., using euclidean distance in the harmonized latent space) should be balanced across the different datasets. For a fixed value of *k*, this property can be evaluated using the entropy of batch mixing [12], which is akin to evaluating a simple *k*-nearest neighbors classifier for the batch identifier (**Online Methods**). While this property is important, it is not sufficient, since it can be achieved by simply randomizing the data. Therefore, in our evaluations we also consider the extent to which the harmonized data retains the original structure observed at each dataset in isolation. Here, we expect that the set of *k*-nearest neighbors of any given cell in its original dataset should remain sufficiently close to that cell after harmonization. This property can be evaluated using a measure we call *k*-nearest neighbors purity (**Online Methods**). Furthermore, because the conservation of *k*-nearest neighbors might be more indicative of a local stability of the algorithm and not of actually preserving global aspects of the datasets, we introduced an additional metric, evaluating the conservation of cluster assignments before and after harmonization. Clearly, these two criteria can be simply optimized by not changing the input datasets, which will result in poor performance with respect to our first measure. Our evaluation therefore relies on both types of measures, namely mixing of data sets and retainment of the original structure. Since our results depend on the neighborhood size *k*, we consider a range of values - from a high resolution (*k* = 10) to a coarse (*k* = 500) view of the data.

We compare scVI to several methods, including MNN [12], Seurat Alignment [13], ComBat [15] and principal component analysis (PCA). In addition, and in order to compare our methods to unpaired data integration approaches based on generative adversarial networks [46], we also tested MAGAN [47]. However, even after manual tuning of the learning rate hyperparameter, the input datasets remain largely unmixed (Supplementary Figure 3). This might be due to the fact that MAGAN was not directly applied to harmonize pairs of scRNA-seq datasets and need more tuning to be applicable in that context. For each algorithm and pair of datasets, we report embeddings computed via a Uniform Manifold Approximation and Projection (UMAP) [48] (Figure 2a, Supplementary Figure 4 – 7) as well as the three evaluation metrics (Figure 2b-c, Supplementary Table 2). Overall, we observed that scVI performs well in terms of mixing, while comparing favorably to the other methods in terms of retainment of the original structure, for a wide range of neighborhood sizes and across all dataset pairs. Reassuringly, our positive results for preservation of the output of a clustering algorithm indicates that scVI and scANVI are also stable with regards to more general aspects of the data. As an illustrative example, consider the results of applying scVI and Seurat Alignment to the Tabula Muris bone marrow datasets (Figure 2a).

**Figure 2:**
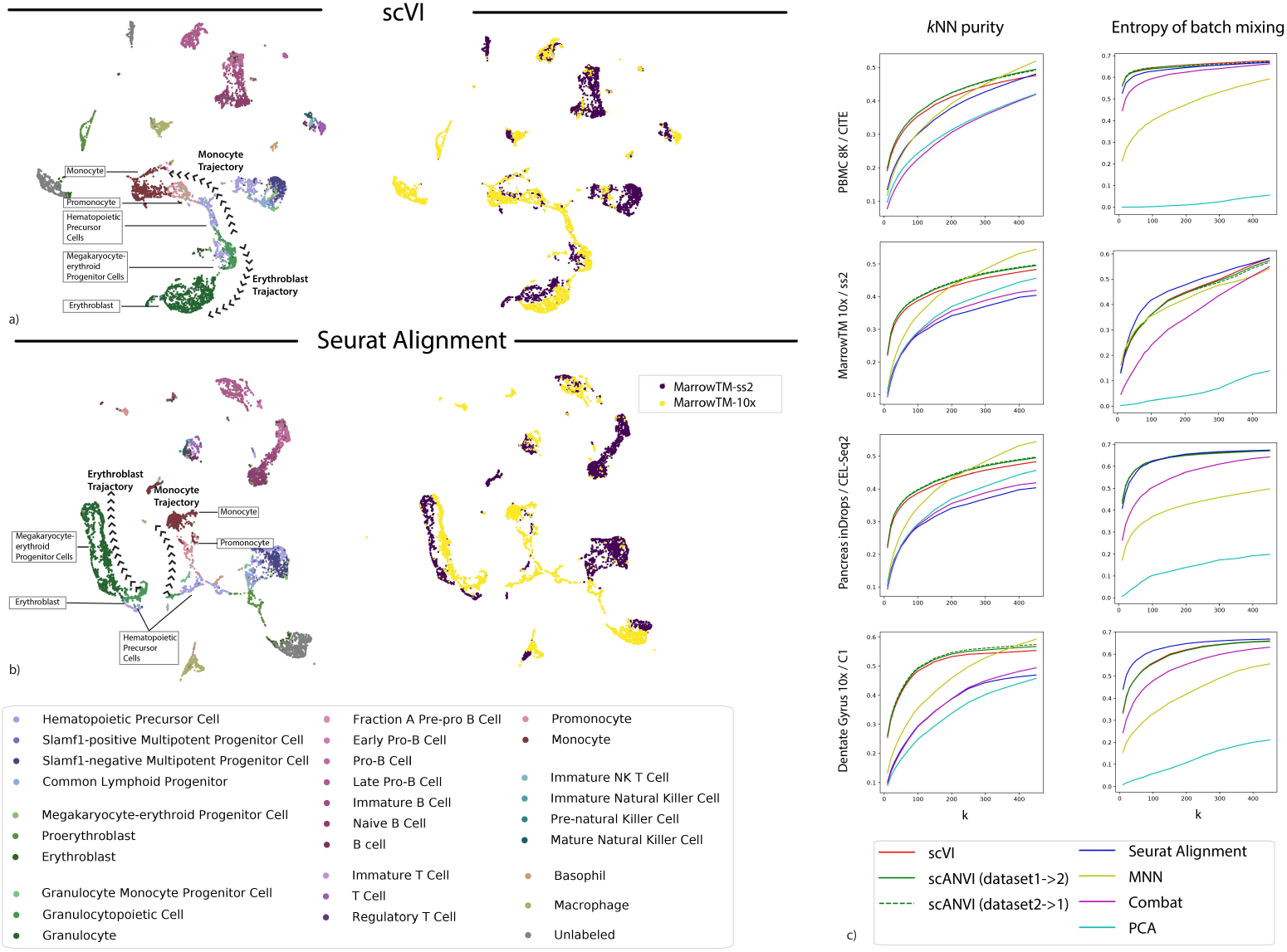
Benchmarking of scRNA-seq harmonization algorithms (*a*) Visualization of the latent space of scVI using Uniform Manifold Approximation and Projection (UMAP [48]) for the bone marrow dataset pair [9], colored by cell type and batches. (*b*) Visualization of the Seurat Alignment latent space of the same dataset. (*c*) Quantitative comparison of the performance of scVI and Seurat Alignment by two measures. On the left retainment of structure for the benchmark algorithms on the four pairs of dataset evaluated via the *k*-nearest neighbors purity, defined as a jaccard index of neighbor sets (higher is better, **Online Methods**). On the right dataset mixing for the benchmark algorithms on the four pairs of datasets evaluated by entropy of batch mixing (higher is better).

While scANVI was designed for the problem of cell state annotation, we also wanted to evaluate its ability to harmonize datasets, which can be seen as a prerequisite. To evaluate this, we consider each dataset pair twice, each time using labels from one of the datasets (exploiting the semi-supervision framework of scANVI) and leaving the other one unlabeled. Reassuringly, we found that scANVI is capable of effectively harmonizing the datasets, with a similar performance to that of scVI in terms of entropy of batch mixing and retainment of the original structure (Figure 2b-c, and Supplementary Table 2). We further explore the performance of scANVI in the annotation problem in the subsequent sections.

### Harmonizing datasets with a different composition of cell types

One of the primary challenges of the harmonization problem is handling cases in which the cell types present in the input datasets only partially overlap or do no overlap at all. Since this is a plausible scenario in many applications, it is important to account for it and avoid over-normalizing or “forcing” distinct cell populations onto each other. To evaluate this, we performed several stress tests in which we artificially manipulated the composition of cell types in the input datasets prior to harmonization. As our benchmark method we use Seurat Alignment, which performed better than the remaining benchmark methods in our first round of evaluation (Figure 2).

As a case study, we used a pair of PBMC datasets (PBMC-CITE [25], PBMC-8K [37]) that initially contained a similar composition of immune cell types (Supplementary Table 3). We were first interested in the case of no biological overlap (Figure 3a-d). To test this, for a given cell type *c*_0_ (e.g., natural killer cells), we only keep cells of this type in the PBMC-CITE dataset and remove all cells of this type from the PBMC-8K dataset. In Figure 3a-b, we show an example of UMAP visualization of the harmonized data, with natural killer cells as the left out cell type *c*_0_. Evidently, when harmonizing the two perturbed datasets with scVI, the natural killer cells appear as a separate cluster and are not wrongly mixed with cells of different types from the other dataset. Conversely, we see a larger extent of mixing in the latent space inferred by Seurat Alignment. A more formal evaluation is provided in Figure 3c-d, which presents our harmonization performance metrics for each cell type averaged across all perturbations (in each perturbation, *c*_0_ is set to a different cell type). We also included scANVI with the true number of cell types (*C* = 6) in this analysis, using the cell labels from the PBMC-CITE dataset.

**Figure 3:**
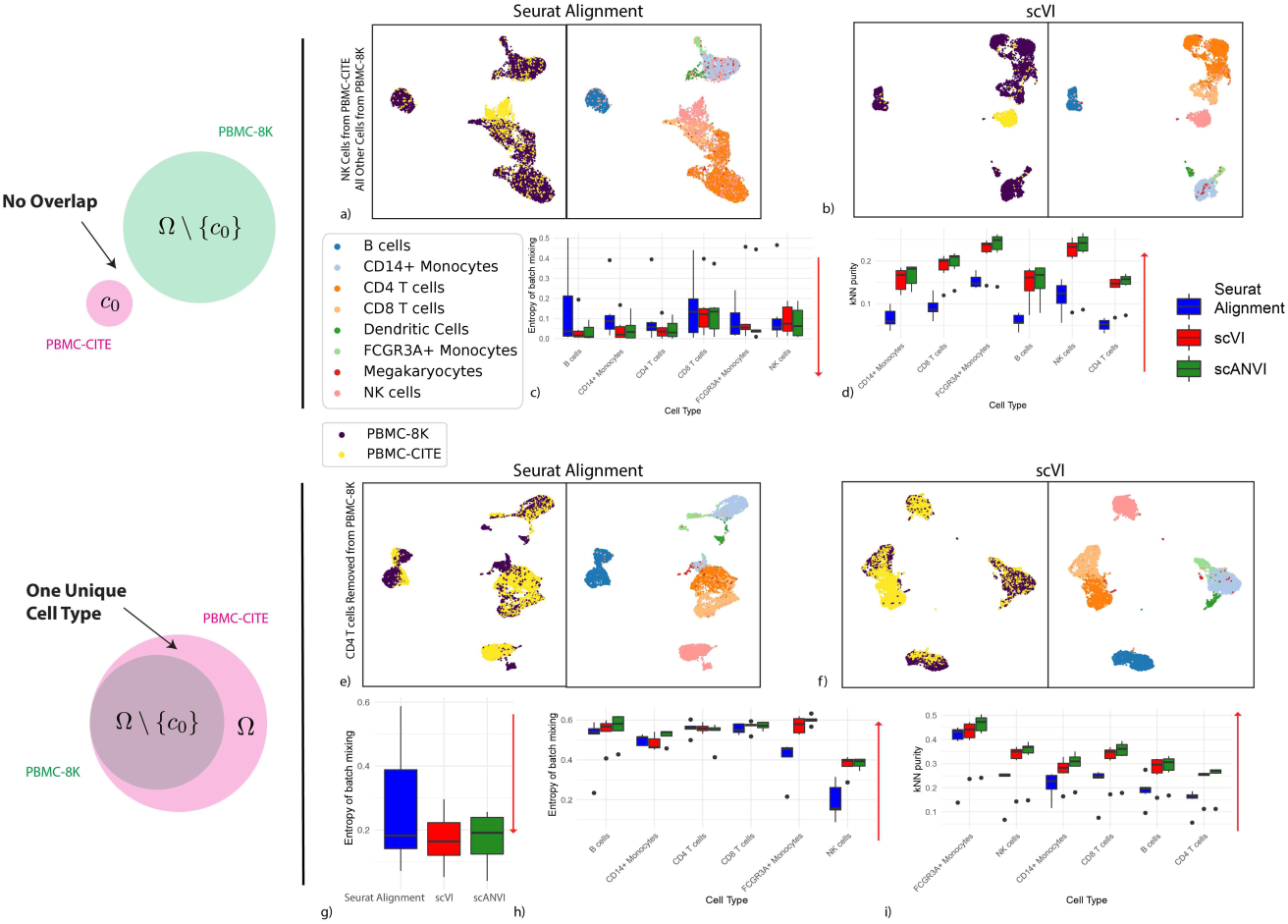
Harmonizing datasets with different cellular composition. (*a* − *d*) show the case when no cell type is shared, where PBMC-8K contains all cells other than cell type *c*_0_ and PBMC-CITE contains only cell type *c*_0_. (*a* − *b*) UMAP visualization for the case where *c*_0_ corresponds to natural killer cells. (*c* − *d*) Harmonization statistics, aggregating the six experiments (setting *c*_0_ to a different cell type in each experiment). (*e* − *i*) show the results of cell type *c*_0_ is removed from the PBMC-8K dataset, while the PBMC-CITE dataset remains the same. (*e* − *f*) UMAP visualization for the case where *c*_0_ corresponds to CD4+ T cells. (*g*) batch entropy mixing for the cell type that was removed. (*h*) batch entropy mixing for each cell type in the 5 experiments where they were not removed. (*i*) Retainment of the original structure, aggregating all 6 experiments. Red arrows indicate the desired direction for each performance measure (e.g., low batch entropy is desirable in (*d*)). The boxplots are standard Tukey boxplots where the box is delineated by the first and third quartile and the whisker lines are the first and third quartile plus minus 1.5 times the box height. The dots are outliers that fall above or below the whisker lines.

Under the ideal scenario of a successful harmonization, we expect both a low entropy of batch mixing (since the datasets do not overlap), and retainment of the original structure. Evidently, both scVI and scANVI exhibit a consistently low level of batch mixing that is better or comparable to that of Seurat Alignment, while retaining the original structure more accurately.

As an additional scenario, we investigated the case where the input datasets contain a similar set of cell types, with the exception of one cell type that appears in only one of the datasets. To simulate this, for a given cell type *c*_0_, we removed cells of this type from the PBMC-8K dataset, and then harmonize the remaining cells with the unaltered PBMC-CITE (which still contains *c*_0_). We show an example of UMAP visualization in Figure 3e-f, removing CD4+ T cells from the PBMC-8K dataset. Evidently, in the scVI latent space, the PBMC-CITE “unique” CD4+ T cell population is not wrongly mixed with cells from the perturbed PBMC-8K dataset, but rather appears as a distinct cluster. For a more formal analysis, Figure 3g-i shows the harmonization statistics for perturbing the six major cell types present in the PBMC datasets. As above, we also evaluated scANVI in this context, using the labels from the unperturbed (PBMC-CITE) dataset.

Figure 3g shows that the entropy of batch mixing from the “unique” populations (averaging over all six perturbations) is low in all three methods (scVI, scANVI and Seurat Alignment), with a slight advantage for scVI and scANVI. Figure 3h-i shows the harmonization statistics for each population, averaging over all shared cell types between the two datasets. Evidently, for the populations that are indeed common to the two datasets, scVI and scANVI are capable of mixing them properly, while preserving the original structure, comparing favorably to Seurat Alignment on both measures. Overall, the results of this analysis demonstrate that scVI and scANVI are capable of harmonizing datasets with very different compositions, while not forcing erroneous mixing. These results are consistent with the design of scVI and scANVI, which aim to maximize the likelihood of a joint generative model, without making *a priori* assumptions about the similarity in the composition of the input datasets.

In a similar but more complex experiment, we also study the case when the two datasets both have their own unique cell types but also share several common cell types. Populations unique to each dataset have low mixing (Supplementary Figure 8a), especially with scVI and scANVI. Conversely, the shared populations have a substantially higher mixing rate (Supplementary Figure 8c).Specifically, scANVI and scVI both mixes shared populations better than Seurat, with a better overall performance for scANVI. Finally, the preservation of original structure is higher scVI and scANVI when compared to Seurat across all cell types, especially for B cells, NK cells and FCGR3A+ Monocytes (Supplementary Figure 8b). Overall, these results demonstrate that our methods do not tend to force wrong alignment of non-overlapping parts of the input datasets.

### Harmonizing continuous trajectories

While so far we considered datasets that have a clear stratification of cells into discrete subpopulations, a conceptually more challenging case is harmonizing datasets in which the major source of variation forms a continuum, which inherently calls for accuracy at a higher level of resolution.

To explore this, we use a pair of datasets that provide a snapshot of hematopoiesis in mice (HEMATO-Tusi [49], HEMATO-Paul [50]; Figure 4). These datasets consist of cells along the transition from common myeloid progenitor cells (Figure 4a-b; middle) through two primary differentiation trajectories myeloblast (top) and erythroblast-megakaryocyte (bottom). Notably, the HEMATO-Tusi dataset contains cells that appear to be more terminally differentiated, which are located at the extremes of the two primary branches. This can be discerned by the expression of marker genes (Figure 4e). For instance the HEMATO-Tusi unique erythroid cell population expresses *Hba-a2* (hemoglobin subunit) and *Alas2* (erythroid-specific mitochondrial 5-aminolevulinate synthase) that are known to be present in reticulocytes [51, 52]. At the other end, the granulocyte subset that is captured only by HEMATO-Tusi expresses *Itgam* and *S100a8*. *S100a8* is a neutrophil specific gene predicted by Nano-dissection [53] and is associated with GO processes such as leukocyte migration associated with inflammation and neutrophil aggregation. *Itgam* is not expressed in granulocyte-monocyte progenitor cells but is highly expressed in mature monocytes, mature eosinophils and macrophages [54]. We therefore do not expect mixing to take place along the entire trajectory. To account for this, we evaluated the extent of batch entropy mixing at different points along the harmonized developmental trajectory. As expected, we find that in most areas of the trajectory the two datasets are well mixed, while at the extremes, the entropy reduces significantly, using either scVI or Seurat Alignment (Figure 4c). Overall, we observe that scVI compares well in terms of both mixing the differentiation trajectories in each dataset and preserving their original, continuous, structure (Figure 4a-d).

**Figure 4:**
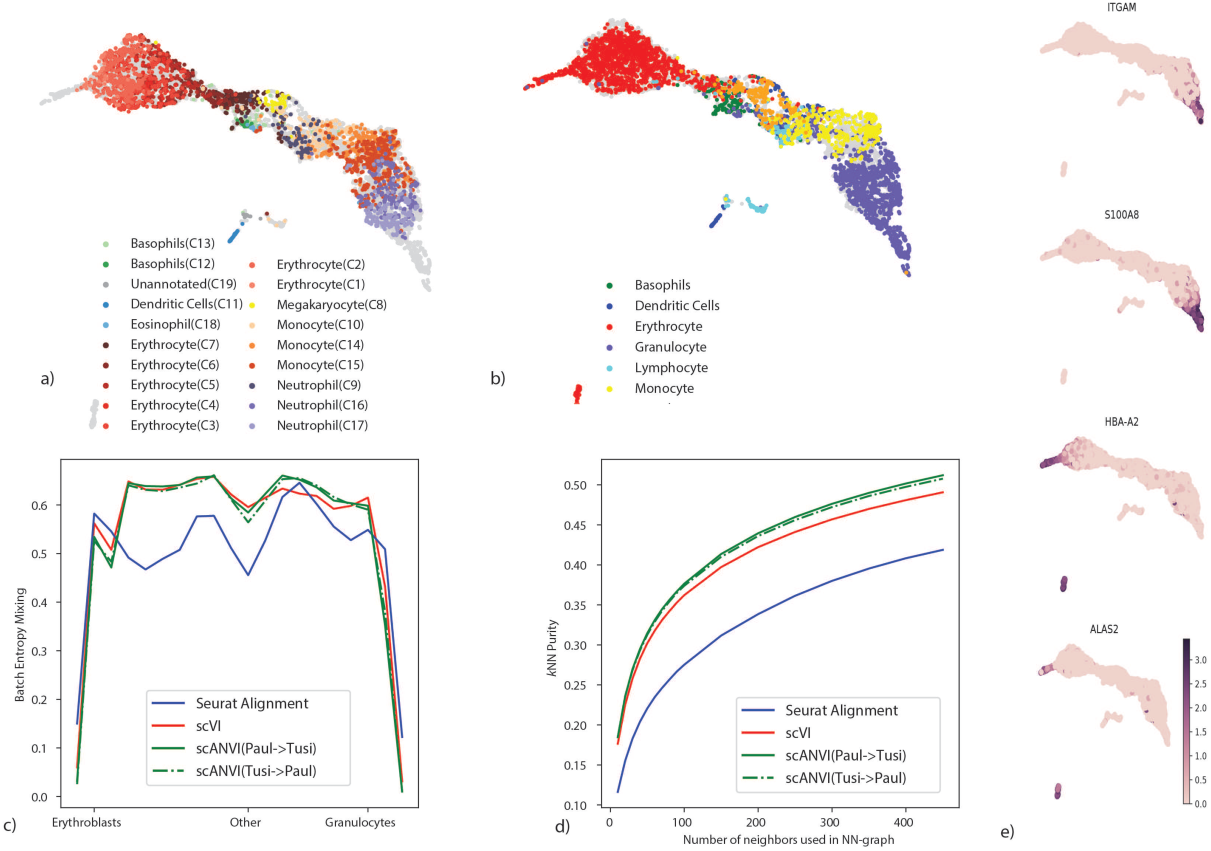
Harmonizing developmental trajectories. (*a* − *b*) UMAP visualization of the scVI latent space, with cells colored by the original labels from either the HEMATO-Paul (*a*) or HEMATO-Tusi (*b*) studies. The cells from the other dataset are colored in gray. (*c*) Batch entropy mixing of 20 of bins HEMATO-Tusi cells ordered by the potential of each cell derived from the PBA algorithm used in the original publication. Potential is a pseudotime measure that describes how differentiated a cell is. (center: common myeloid progenitors; moving left: erythrocyte branch; moving right: granulocyte branch). (*d*) *k*-nearest neighbors purity for scVI, Seurat, and scANVI. (*e*) Expression of marker genes that help determine the identity of batch-unique cells.

To validate scANVI in this context as well, we provided it with the categorical labels of cells along the two developmental trajectories, indicating their cell state (Figure 4c-d and Supplementary Figure 9). Even though this labeling scheme does not explicitly account for the ordering between states, we observe that scANVI is capable of mixing the two datasets, while retaining their original structure, achieving a level of accuracy comparable to that of scVI and better than that of Seurat Alignment.

### Rapid integration of multiple datasets

To demonstrate the scalability of our framework in the context of harmonizing multiple (and possibly large) dataset, we ran scVI to integrate a cohort of 26 datasets spanning 105,476 cells from multiple tissues and technologies, which was made available by the authors of Scanorama (a method based on truncated singular value decomposition followed by nearest neighbor matching [19]). Using the hardware specified in the original paper [19] (Intel Xeon E5-2650v3 CPU limited to 10 cores with 384 GB of RAM), Seurat Alignment and MNN required over 24 hours, while Scanorama completed its run in 20 minutes. Using a simpler configuration (eight-core Intel i7-6820HQ CPU with 32 GB RAM) along with one NVIDIA Tesla K80 GPU (GK210GL; addressing 24 GB RAM), we found that scVI integrates all datasets and learns a common embedding in less than 50 minutes. This running time is competitive considering the reduced memory availability and the increased complexity of our model, compared to that of Scanorama. Notably, all the downstream analyses, such as annotation, differential expression or visualization can be operated by accessing the latent space or via forward passes through the neural networks. Since these access operations can be conducted very efficiently [29], the dominant factor, on which we focused our run time analysis, is the time required for model fitting. Considering the results, the latent space of scVI recapitulates well the major tissues and cell types (Supplementary Figure 10), and the position of cells in the latent space provides an effective predictor for the cell type label (Supplementary Figure 10 and **Online Methods**).

### Transferring cell type annotations between datasets

We next turned to evaluate scVI and scANVI in the context of harmonization-based annotation. Here, we test the extent to which annotations from a previously annotated dataset can be used to automatically derive annotations in a new unannotated dataset. For scVI and Seurat Alignment, we derive the annotations by first harmonizing the input datasets and then running a *k*-nearest neighbors classifier (setting *k* to 10) on the joint latent space, using the annotated cells to assign labels to the unannotated ones. Conversely, scANVI harmonizes the input datasets while using any amount of available labels. The prediction of unobserved labels is then conducted using the approximate posterior assignments *q*_Φ_(*c* | *x*) of cell types (**Online Methods**), which can be derived directly from the model. An alternative approach, which we also include in our benchmark was taken by scmap-cluster [28], which instead of harmonizing, directly builds a classifier based on the labeled cells and then applies this classifier to the unlabeled cells. Finally, we also applied the domain adaptation method Correlation Alignment for Unsupervised Domain Adaptation (CORAL, [55]). This method was not initially developed for single-cell analysis but is an insightful benchmark from the machine learning literature.

We start by exploring the four dataset pairs in Figure 2, which have been annotated in their respective studies. In each experiment, we “hide” the cell type annotations from one dataset and transfer the second dataset labels to the first one. As a measure of performance, we report the weighted accuracy, which is the percent of cells that were correctly assigned to their correct (hidden) label, averaging over all labels (**Online Methods**). Importantly, the annotations in this first set of case studies were derived computationally. For example, by first clustering the cells, looking for marker genes expressed by each cluster and then assigning labels to the clusters accordingly. This level of annotation therefore makes the prediction problem relatively easy, and indeed, while we find that overall scANVI predicts unobserved labels more accurately, the differences between the methods are mild (Supplementary Figures 11 and 12). Notably, CORAL achieves overall competitive performance except when transferring labels on the MarrowTM pairs, from 10x to Smart-Seq2. In this specific instance, CORAL maps most of the cells to a single label (incidentally, while this label marks cells that are transcriptionally similar, it is defined by the authors as an unknown class “NA”, corresponding to cells that cannot be confidently assigned or low quality cells according to the authors of [9]), which might be due to its linear transformation of the feature space.

To evaluate the accuracy of annotations without the need for computationally-derived labels, we turned to the PBMC-CITE dataset which includes measurements of ten key marker proteins in addition to mRNA [25], and the PBMC-sorted dataset [3], where cells were collected from bead purifications for eleven cell types (Supplementary Table 4). We applied scVI and scANVI to harmonize and annotate these two datasets along with a third dataset of PBMC (PBMC-68K [3]). Our analysis contains a combined set of *n* = 169, 850 cells from the three datasets altogether. To generate a realistic scenario of cell type annotation, we only provide access to the experimentally-based labels from the PBMC-sorted dataset (Figure 5a-c). As an additional benchmark, we also evaluate Seurat Alignment, which was tested after removal of a randomly selected subset (40%) of the two large datasets (PBMC-68K and PBMC-sorted) due to scalability issues. Considering our harmonization performance measures (i.e., retainment of the original structure and batch mixing), we observe as before that scVI and scANVI perform similarly and compare favorably to Seurat Alignment. We then evaluated the accuracy of assigning unobserved labels, focusing on the PBMC-CITE dataset. Instead of using the labels from the original PBMC-CITE study as ground truth (which were computationally derived), we used the protein data, which provides an experimentally-derived proxy for cell state. To this end, we quantified the extent to which the similarity between cells in the harmonized mRNA-based latent space is consistent with their similarity at the protein level (**Online Methods**). We first computed the average discrepancy (sum of squared differences) between the protein measurements in each cell and the average over its *k*-nearest neighbors. As a second measure we computed for each PBMC-CITE cell the overlap between its *k*-nearest PBMC-CITE neighbors in the harmonized mRNA-based space and in the protein space. We then report the average across all cells in Supplementary Figure 13. Evidently, scANVI outperformed both scVI and Seurat Alignment for a wide range of neighborhood sizes, providing a representation for the mRNA data that is more consistent with the protein data (Figure 5c).

**Figure 5:**
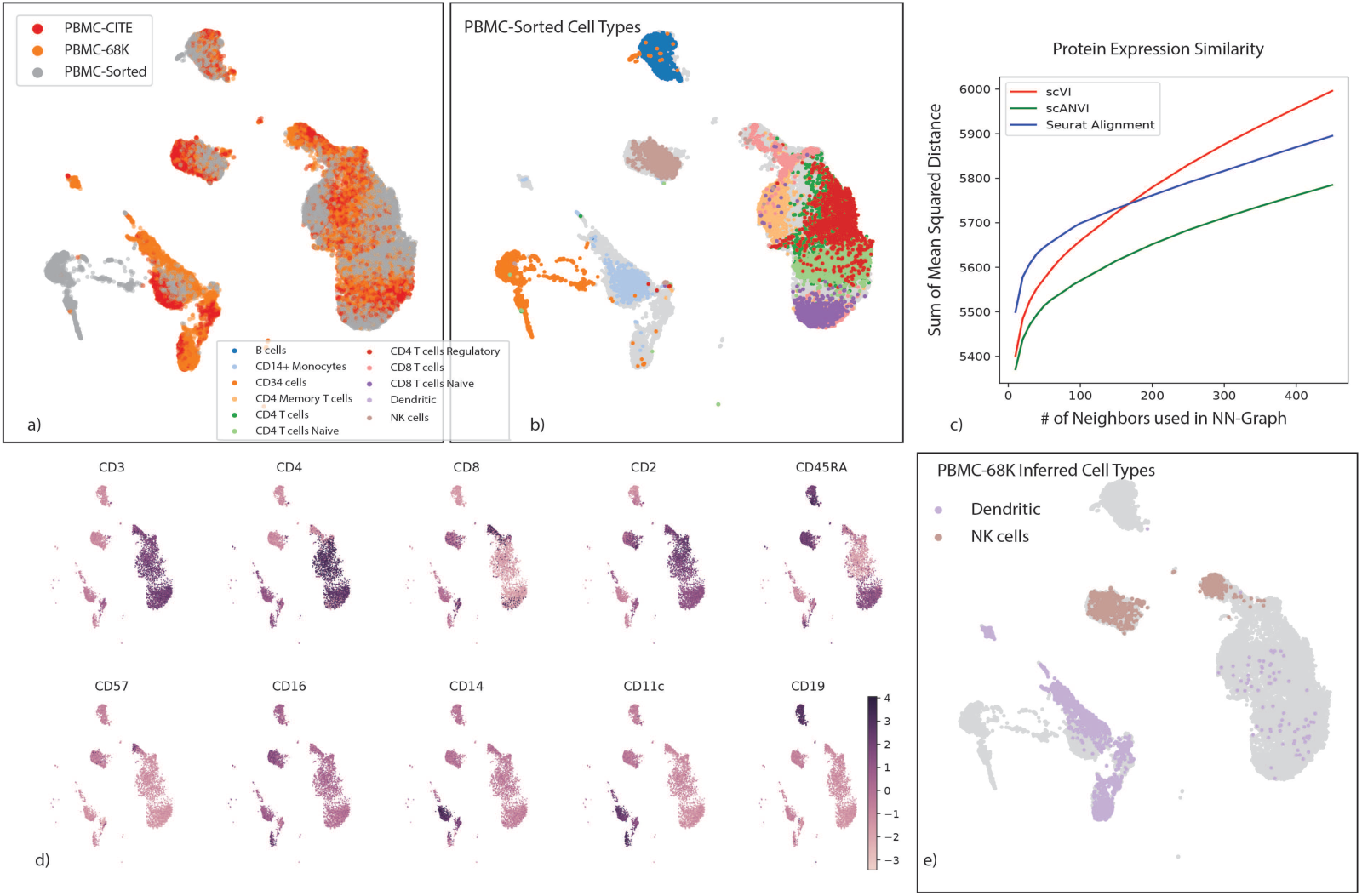
Validation of cell type annotations using additional metadata. (a-b) UMAP plot of the scANVI latent space inferred for three harmonized datasets: PBMC-CITE [25], PBMC-sorted [3], and PBMC-68K [3]. Cells are colored by the batch they come from (*a*) and the PBMC-sorted labels(*b*). Cells from the PBMC-Cite and PBMC-68K are colored gray in (*b*). (c) The consistency of the harmonized PBMC-CITE mRNA data with the respective protein measurements, evaluated by mean squared error and for different neighborhood size. Lower values indicate higher consistency. (*d*) UMAP plot of the scANVI latent space, where cells are colored by normalized protein measurement. Only PBMC-CITE cells are displayed. (*e*) UMAP plot of the scANVI latent space, with cells from the PBMC-68k dataset colored according to their original label. For clarity of presentation, only cells originally labeled as dendritic cells or natural killer cells are colored. Evidently, a large number of these cells are mapped to a cluster of T-cells (right side of the plot).

As further support for the validity of these results, we confirmed that the labels inferred by scANVI in the PBMC-CITE dataset are consistent with the protein expression values (e.g., observing a uniquely high level of *CD19* expression in cells assigned with the “B cell” label; Figure 5d and Supplementary Figure 13). Interestingly, when considering the latent space learned by scANVI, we observed a certain amount of possible mislabeling in the original study of the PBMC-68K dataset (Figure 5e). In that study, the PBMC-68K cells were assigned with labels by taking the maximum correlation with cell subsets from the experimentally-annotated PBMC-sorted data. However this approach might be error prone, most likely due to low sensitivity and the influence of genes that are less relevant to cell type classification. Specifically, we observe a substantial cells that are labeled by scANVI as T cells, but originally labeled as dendritic cells or natural killer cells. This re-annotation as T cells is supported by the expression of marker genes [56] of the respective cell subsets (Supplementary Figure 13).

### Cell type annotation in a single dataset based on “seed” labels

An important variant of the annotation problem lies within the context of an *ab initio* labeling of a single dataset where only a subset of the cells can be confidently annotated based on the raw data. This increasingly prevalent scenario may result from limited sensitivity of the scRNA-seq assay, where marker genes may only be confidently observed in a small subset of cells. One common way to address this problem is to compute some form of a distance metric between cells (e.g., after embedding with scVI or using Seurat PCA) and then assign labels based on proximity to annotated cells [3]. To benchmark our methods, we consider two such predictors: the first is clustering the cells and taking a majority vote inside each cluster, and the second is taking the majority vote of the *k*-nearest neighbors around each unannotated cell (*k* = 10). While these approaches are quite straightforward, their accuracy might suffer when the data does not form clear clusters [49], or when differences between labels are too subtle to be captured clearly by a transcriptome-wide similarity measure. To address these issues, scANVI takes an alternative approach, namely learning a latent embedding that is guided by the available labels, and then producing posterior probabilities for assigning labels to each cell.

As a case study, we compiled a dataset consisting of several experimentally sorted and labeled subsets of T cells from the PBMC-sorted dataset, including CD4 memory, CD4 naive, CD4 regulatory and CD8 naive. To make our analysis more realistic, we assume that the labels are completely unknown to us and therefore begin by trying to assign each T cell to its respective subset using marker genes (12 altogether; see **Online Methods**). Notably, several important biomarkers (*CD4*, *CTLA4*, and *GITR*) are detected in less than 5% of the cells, which renders their use for annotation not straightforward. Furthehrmore, many of these biomarkers are sparsely expressed to the extent that they are likely to be filtered out in the gene selection step of most harmonization procedures (Figure 6a).

**Figure 6:**
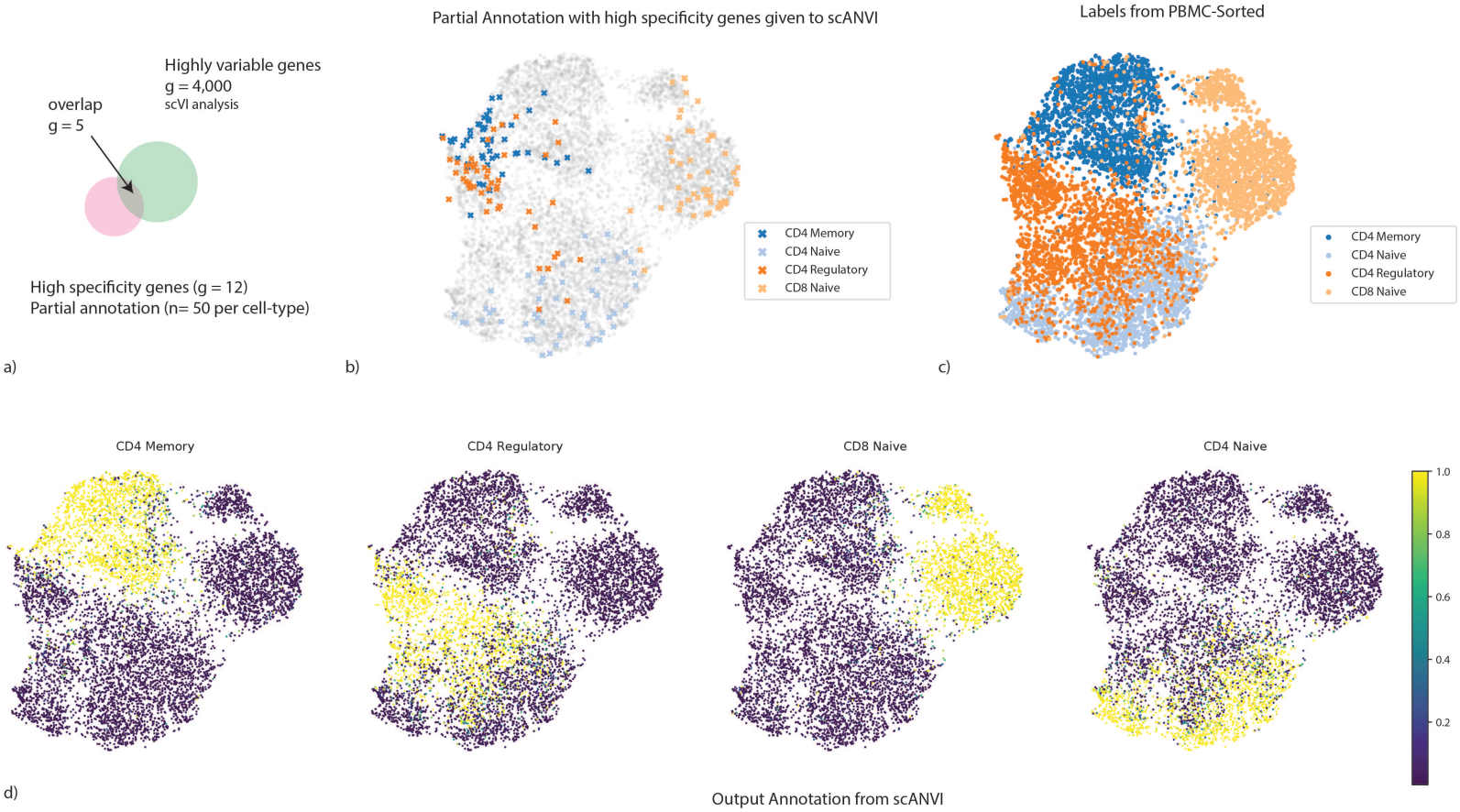
Cell type annotation in a single dataset using “seed” labeling. (*a*) discrepancies between marker genes that can be used to confidently label cells and highly variable genes in scRNA-seq analysis. (*b* − *d*) UMAP plot of the scVI latent space. (*b*) Seed cells are colored by their annotation (using known marker genes). (*c*) PBMC-sorted cell type labels from the original study based on marker-based sorting (*d*) The posterior probability of each cell being one of the four T cell subtypes obtained with scANVI.

To analyze this dataset, we first computed a signature score for each cell and for each label (i.e., T cell subset) using the scaled raw expression values of the respective marker genes (**Online Methods**). We then designated the top 50 scoring cells in each subset as the seed set of cells that are confidently annotated for that subset (Figure 6b). Reassuringly, this partial annotation is in agreement with the experimentally derived cell type labels available for this dataset (Figure 6c). However, this dataset does not form clear clusters, and in particular the seed sets of cells are not well separated. Such an observation makes clustering-based approaches potentially less precise. Indeed, using *k*-means clustering on the scVI and Seurat PCA latent space, we find that 74% and 72% of the cells were assigned with their correct label. Similar analysis with two additional popular clustering algorithms (DBSCAN [57] and PhenoGraph [58]) further emphasizes the challenge of a cluster-based approach on this data. Specifically, DBSCAN does not partition the data into more than one cluster (scanning through a large number of parameter values; **Online Methods**), and PhenoGraph predicts 9 clusters and achieves an accuracy of 41% (Supplementary Figure 14).

Consistent with these results, the application of a *k*-nearest neighbors classifier resulted in a similar level of accuracy in the Seurat PCA latent space (71%), which is slightly improved when replacing it with the scVI latent space (73%; Supplementary Figure 14). Conversely, after fitting the scANVI model based on this partial labeling, the annotation posterior *qΦ*(*c* | *z*) (Figure 6d) provides a substantially more accurate cell type assignment, with 84% of cells annotated correctly.

While scANVI has been designed to handle discrete (but not continuous) labels, we hypothesized that gradual transition between cell states may still be captured by the uncertainty of label assignment. We tested it using simulated data [59] that consists of a set of “end-point” states along with intermediary states that connect them (**Online Methods**, Supplementary Figure 15a). We provided labels only to end-point cells, and investigated the label assignment scores calculated for the intermediary cells. We find that scANVI provides a range of assignment probability values and that these values are proportional to the distance from the respective end points (Supplementary Figure 15b-g). Conversely, the scores provided by scmap tend to be more extreme (Supplementary Figure 15h-i), thus less reflecting the continuous nature of the data.

### Cell type taxonomy and hierarchical classification with scANVI

Another subtle yet important variation of the annotation problem is when the labels are not mutually exclusive but rather form a taxonomy of cell types or states. To effectively annotate cells in this setting, we extended scANVI to perform hierarchical classification, which as before we carry out from first principles, relying on probabilistic graphical models (**Online Methods**). To demonstrate this extended version, we use a dataset of the mouse nervous system [60] that was annotated using a cell type taxonomy with several levels of granularity. At the lowest (most granular) level, the cells are stratified into 265 cell sub-types. At the second lowest level of granularity these 265 subtypes are grouped into 39 subsets, each corresponding to a more coarse definition of a cell type.

We evaluate the ability of scANVI as well as the competing methods at inferring the most granular level of labels when provided with partial “seed” annotation — namely label information for 5 randomly selected cells per label (which accounts for an overall of 0.8% of the cells). We first observe that Seurat PCA followed by a *k*-nearest neighbors classifier provides a weighted accuracy of 23% (averaging over all cell types). While this might seem like a low accuracy, it is in fact far from trivial since the expected weighted accuracy of a random classifier or a constant predictor is of around 1*/*265 ⇡ 0.3%. Such low numbers are due to the high number of labels at this highly granular scale. scVI provides a substantially better, yet still lower level of accuracy at 32%. Interestingly, when scANVI is used without accounting for hierarchy, its performance is similar to the unsupervised scVI (at 32%), which might result from very large number of labels that may require hyperparameter tuning (e.g., increasing the number of classifier training epochs; see **Supplementary Note 3**). However, when we take the hierarchy of the labels into account, the performance of scANVI increases to 37%, thus outperforming the other methods by a significant margin. Notably, while we tested the extrapolation of seed labeling and the hierarchical mode only in the context of a single dataset, this variation of the scANVI model can also be directly applied in the context of multiple datasets (i.e., transferring hierarchical annotations between datasets).

### Hypotheses testing in harmonized datasets: the case of differential expression

With their probabilistic representation of the data, scVI and scANVI each provide a natural way of performing various types of hypotheses testing (**Online Methods**). This is different from other approaches [12, 13, 18, 19, 20] where the dataset alignment procedures do not carry direct probabilistic interpretation, and the resulting harmonized data can thus not be directly used for these purposes.

To demonstrate this, we focus on the problem of differential expression. As a first case study, we use two of the PBMC datasets (PBMC-8K and PBMC-68K) and looked for differentially expressed genes in two settings: comparing the B cells to dendritic cells, and similarly for CD4+ versus CD8+ T cells. For evaluation, we used reference sets of differentially expressed genes that were obtained from published bulk-level analysis of similar cell subsets (microarrays, [61, 62], as in [29]). While this benchmark relies on real data, a clear caveat is the lack of a well defined ground truth. To address this, we used a second benchmark based on simulations with Symsim [59]. The simulated data consists of five subpopulations of varying degrees of transcriptional distance, profiled in two different “batches” of different technical quality (**Online Methods**, Supplementary Figure 16). This framework allowed us to derive an exact log fold changes (LFC) between every pair of simulated subpopulations, which enable a more accurate evaluation of performance.

In both benchmark studies, we assume that labels are only available for one of the two input batches or datasets (in the real data we assume that PBMC-8K is the annotated one). To apply scVI, we first harmonized the input pair of datasets and transferred labels using a *k*-nearest neighbors classifier on the joint latent space (*k* = 10). We then consider these annotations (predicted and pre-labeled) as fixed and sample 100 cell pairs, each pair consisting of one cell from each population. For each cell pair we sample gene expression values from the variational posterior, while marginalizing over the different datasets, to compute the probability for differential expression in a dataset-agnostic manner. Aggregating across all selected pairs results in approximate Bayes factors that reflect the evaluated extent of differential expression (**Online Methods**). Since scANVI assigns posterior probability for associating any cell to any label, it enables a more refined scheme. Specifically, instead of sampling pairs of cells we are sampling pairs of points in the latent space, while conditioning on the respective label. This approach therefore does not assume a fixed label for each cell (or point in latent space) as in the scVI scheme, but rather a distribution of possible labels thus making it potentially more robust to mis-labeling. As reference, we also included edgeR [63] using the same labels as scVI. Notably, edgeR was shown to perform well on scRNA-seq data [64] and uses a log-linear model to control for technical sample-to-sample variation.

To evaluate performance on the real data, we defined genes as differentially expressed if the adjusted p-value in the reference bulk data (provided by [61, 62]) was under 5%. Considering these genes as positive instances, we calculated the area under the ROC curve (AUROC) based on rank ordering the inferred Bayes factors (for scVI and scANVI) or p-values (for edgeR). Since the definition of positives genes required a somewhat arbitrary threshold, we also used a second score that evaluates the reproducibility of gene ranking (bulk reference vs. single-cell; considering all genes), using the Kendall rank correlation coefficient (Supplementary Figure 17c). As a reference, we look at the accuracy of differential expression analysis in each PBMC dataset separately (using their prior annotations to define the sets of cells we are comparing), which can computed with scVI (as in [29]) and edgeR. Reassuringly, we observe that the performance of scVI on the joint data is not lower than it is in either dataset in isolation. We also find that while scVI performs moderately better than scANVI, both methods compare favorably to edgeR in their accuracy.

In our simulations, we considered differential expression between every possible pair out of the five simulated subpopulations. For evaluation, we computed the Spearman and Kendall rank correlation coefficients between the true LFC and the inferred Bayes factors (for scVI and scANVI) or estimated LFC (for edgeR). Our results in Supplementary Figure 17ab show that with this artificial, yet more clearly defined objective, scVI was substantially more accurate than edgeR and that in the harmonized data scANVI provided more exact and stable estimates than scVI (Supplementary Figure 17).

Mislabeling of a certain proportion of cells in a dataset is a plausible scenario that may occur in any study. An important challenge is therefore to maintain the validity of downstream analysis despite such “upstream” annotation errors. To evaluate robustness in this setting, we repeated the simulation analysis, while introducing labeling errors at different rates. Specifically, prior to evaluating differential expression between two simulated sub-populations, we flip the labels of a certain proportion (up to 30%) of the respective cells in the annotated batch. We then proceed as before and assign labels to cells in the unannotated batch by scVI or scANVI, followed by differential expression analysis. Our results (Supplementary Figure 18a) suggest that scANVI is clearly more robust to this type of mislabeling than scVI (or edgeR, applied on the scVI- derived labels). Repeating the same analysis on the PBMC data (where the differential expression ground truth is obviously not available), we observe similar level of robustness in scANVI, albeit with not much difference compared to scVI and edgeR.

Overall, our results demonstrate that both scVI and scANVI are capable of conducting differential expression effectively, while working directly on a harmonized dataset. Furthermore, we observe that both methods and especially scANVI are robust to mislabeling, providing further motivation for explicitly modeling label uncertainty.

## Discussion

In this study, we demonstrated that scVI provides a principled approach to harmonization of scRNA-seq data through joint probabilistic representation of multiple dataset, while accounting for technical hurdles such as variable library size and limited sensitivity. We have demonstrated that scVI compares favorably to other methods in its accuracy and that it scales well, not only in terms of the number of cells (as in [29]) but also the number of input datasets (as opposed to other methods that work in a pairwise fashion and therefore scale quadratically with dataset size [19]). We have also shown that the harmonization step of scVI provides an effective baseline for automated transfer of cell type labels, from annotated datasets to new ones.

While the performance of scVI in the annotation problem compares favorably to other algorithms, it does not make use of any existing cell state annotations during model training, but rather after the latent space has been learned. To make better use of these annotations (which may be available for only some of the input datasets or only some cells within a dataset), we developed scANVI, a semi-supervised variant of scVI. While the latent space of scVI is defined by a Gaussian vector with diagonal unit variance, scANVI uses a mixture model, which enables it to directly represent the different cell states (each corresponding to a mixture component; see **Online Methods**) and provide a posterior probability of associating each cell with each label. We have demonstrated that similar to scVI, scANVI is capable of harmonizing datasets effectively. In addition, scANVI provides a way to address a number of variants of the annotation problem. Here, we have first shown that it performs well in the most prevalent application of transferring labels from a reference dataset to an unannotated one. We then demonstrated that scANVI can be used in the context of a single unannotated dataset, where high confidence (“seed”) labels are first inferred for a few cells (using marker genes) and then propagated to the remaining cells. Finally, we have shown that scANVI is especially useful in the challenging case where the differences between cell states are too subtle to be captured clearly by a transcriptome-wide similarity measure, as well as in the case where the labels are organized in a hierarchy.

Notably, although scANVI achieves high accuracy when transferring labels from one dataset to another, it was not designed to automatically identify previously unobserved labels. Indeed, in Supplementary Figure 19 we demonstrate that increasing the number of labels in the model (*C*) to values beyond the number of observed labels does not alter the results much. Nevertheless, we observed that unannotated cell populations that have an unobserved label are associated with low levels of mixing between the input datasets. We therefore advocate that clusters from an unannotated dataset that do not mix well should be inspected closely and, if appropriate, should be manually assigned with a new label.

One concern in applying methods based on neural networks [21, 65, 66, 67, 68] in single-cell genomics and other domains is the robustness to hyperparameters choices [69]. This concern has been addressed to some extent by recent progress in the field, proposing search algorithms based on held-out log-likelihood maximization [67]. In this manuscript, we used an alternative approach that is more conducive for direct and easy application of our methods — namely we fix the hyperparameters and achieve state-of-the-art results on a substantial number of datasets and case studies.

The development of scVI and scANVI required several modeling and implementation choices. In **Supplementary Note 4**, Supplementary Figures 20 and 21 we discuss the rationale behind the choice of a zero-inflated negative binomial (ZINB) distribution as well as robustness to choice of priors. Briefly, we find that exclusion of zero inflation from the model results in approximately similar performance, except for the case of harmonizing Smart-Seq2 and 10x data sets, in which ZINB performs significantly better. Such results might suggest that zero-inflation may be more suitable for certain technologies than others. Similarly, we investigate the prior on the library size which is defined per batch and show that computing the same prior for both the datasets (rather than each dataset individually, as we do by default) affects the performance only in the case of harmonizing the same pair of Smart-Seq2 and 10x datasets. Since these datasets have very different sources of technical noise, this may suggest that it is indeed advisable to explicitly account for such differences during model fit.

Another important consideration while designing statistical models is the trade-off between goodness of fit and interpretability, which is still an open topic in machine learning research. Simple models such as the latent Dirichlet allocation [70] might not be particularly suited for scRNA-seq noise but are yet of interest, e.g., due to the immediate interpretation of word-level (i.e., gene-level) contributions to topics (i.e., cell types) [71]. Further effort in the use of deep generative models for applications in computational biology should come with attempts to perform model interpretation. For instance, SAUCIE [21] experimentally proposes to add an entropy regularization to a hidden layer of its denoising auto-encoder in order to infer clustering. Future principled efforts may focus on putting a suitable prior such as sparsity on neural networks weights (e.g., as in [72]). That way, individual neurons of the last hidden layer of the generative model would correspond to individual gene modules, directly readable from the weight sparsity motifs. Finally, as recent preprints propose proof of concepts for integrating single cell data across different data modalities such as Single molecule fluorescent in situ hybridization (smFISH), RNA-seq, ATAC-seq and DNA methylation [18, 20], further work can utilize probabilistic graphical models that quantify measurement uncertainties in each assay, as well as the uncertainties of transferring information between modalities (e.g., predicting unmeasured gene expression in smFISH data as in [73]).

An important distinguishing feature of both scVI and scANVI is that they rely on a fully-probabilistic model, thus providing a way to directly propagate uncertainties to any downstream analysis. While we have demonstrated this for differential expression analysis and cell type annotation, this can be incorporated to other tasks, such as differential abundance of sub-populations in case-control studies, correlation between genes and more. We therefore expect scVI, scANVI and similar tools to be of much interest as the field moves toward the goal of increasing reproducibility and consistency between studies and converging on to a common ontology of cell types. In particular, we expect scANVI to be especially useful for transferring labels while taking into account the uncertainty, or in the case of a more complex label structure such as hierarchical cell types. Both scVI and scANVI are available as an open-source software at https://github.com/YosefLab/scVI. The code for reproducing the results in this manuscript has been deposited at https://doi.org/10.5281/zenodo.2529945.

## Acknowledgments

CX, RL and NY were supported by grant U19 AI090023 from NIH-NIAID. We thank Maxime Langevin, Yining Liu and Jules Samaran for helpful discussions and work on the scVI codebase as well as Allon Wagner and Chao Wang for their help on the choice for high specificity genes in the T cell study.

## Author contributions

RL, EM, JR and NY conceived the statistical model. EM developed the software. CX, RL and EM applied the software to real data analysis. CX, RL, JR, NY, and MIJ wrote the manuscript. NY and MIJ supervised the work.

## Online Methods

### scANVI: an extension to scVI for harmonization through semi-supervised annotation

scVI is a hierarchical Bayesian model [29] for single-cell RNA sequencing data with conditional distributions parametrized by neural networks. The graphical model of scVI (Figure 1c) is designed to disentangle technical signal (i.e., library size discrepancies, batch effects) and biological signal. We propose in this manuscript an extension of the scVI model to include information about cell types in the generative model. We name this extension scANVI (single cell ANnotation using Variational Inference). In our generative model, we assume each cell *n* is an independent realization of the following generative process.

Let *K* be the number of datasets and *C* be the number of cell types across all datasets (including cell types that are not observed). Let c describe the expected proportion of cells for each cell type. As in general this information is not available to the user, we consistently use a non-informative prior **c** = 1/*C* in the manuscript. Although some prior information about proportions of cell type is generally accessible, we observe that using the non-informative prior allows us to recover the correct proportion of cells, and adjustment to the prior is not required. In addition, since in comparative studies such as disease case-control comparisons, or between tissue comparisons of immune cells [74] we might not want to bias the estimate of cell-type proportion by prior knowledge. Latent variable

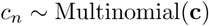

describes the cell type of the cell *n*. Latent variable

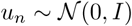

is a low-dimensional random vector describing cell *n* within its cell type. Conceptually, this random variable could describe cell-cycles or sub-cell types. By combining cell type information *c*_*n*_ and random vector *u*_*n*_, we create a new low-dimensional vector

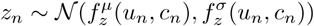

where 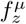 and 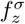 are two functions parametrized by neural networks. Let *s*_*n*_ encode dataset information. Given 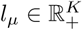 and 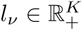 specified per dataset as in [29], latent variable

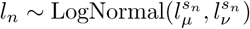

encodes a cell-specific scaling factor. As the prior are adjusted per dataset, our inference procedure will shrink the posteriors towards dataset specific values. This is particularly useful when aligning datasets with dramatically different library size values. Let 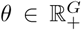 encode a gene specific inverse dispersion parameter (inferred as in [29]). Conditional distribution *x*_*ng*_ | *z*_*n*_, *l*_*n*_, *c*_*n*_, *s*_*n*_ is conform to the one from the scVI model

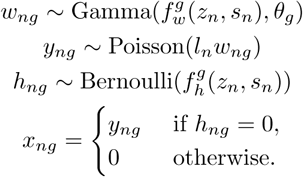

where *f*_*w*_ and *f*_*h*_ are functions parametrized by neural networks. *f*_*w*_ has a final softmax layer to represent normalized expected frequencies of gene expression as in [29]. Let us note that the resulting distribution for the counts is zero-inflated negative binomial. However, it is straightforward using our implementation to use a negative binomial or a Poisson noise model instead. In this model, annotation *c*_*n*_ can be either observed or unobserved following [31, 33], which is useful in our applications where some datasets would come partially labeled or unlabeled. Only the first part of the generative model, as separated above, differs from the original scVI formulation. This corresponds to the top part of the new representation of the graphical model in Figure 1b.

### Hierarchical classification of cells onto a cell type taxonomy

For hierarchical label propagation in scANVI, we propose a extension of the formerly presented model by modifying the variable *c*_*n*_ to be a tuple where each entry denotes the label at a given level of the hierarchy. Our approach is similar to previous work in robustness to noisy labels [75] and hierarchical multi-labels flavors of classification problems [76]. We detail the case for a depth of level two in **Supplementary Note 5** though our approach can in principle be adapted to arbitrary depths.

### Posterior inference

We rely on collapsed variational inference, a standard approximate Bayesian inference procedure that consists in analytically integrating over some of the random variables [77] before optimizing the parameters. As we proved in [29], we can integrate the random variables {*w*_*ng*_, *y*_*ng*_, *h*_*ng*_} to simplify our model at the price of a looser though tractable lower bound (*x*_*ng*_ | *z*_*n*_, *l*_*n*_, *s*_*n*_ is zero-inflated negative binomial). This procedure reduces the number of latent variable and avoid the need for estimating discrete random variables, which is a harder problem. We then use variational inference, neural networks and the stochastic gradients variational Bayes estimator [30] to perform efficient approximate inference over the latent variable {*z*_*n*_, *u*_*n*_, *c*_*n*_, *l*_*n*_}. We assume our variational distribution factorizes as

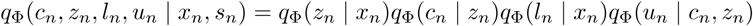

Following [31, 33], we derive two variational lower bounds: one 𝓛 in the case of *c*_*n*_ observed for *pΘ*(*x*_*n*_, *c*_*n*_ | *s*_*n*_) and a second 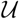 in the case of *c*_*n*_ non-observed for *pΘ*(*x*_*n*_ | *s*_*n*_) where Θ are all the parameters (neural networks and inverse-dispersion parameters). Equations to derive the *evidence lower bound* (ELBO) are derived in **Supplementary Note 6**. We optimize the sum 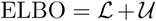 over the neural networks parameters and the inverse-dispersion parameters (in a variational Bayesian inference fashion). Remarkably, the approximate posterior *qΦ*(*c*_*n*_ | *z*_*n*_) can be used as a classifier, assigning cells to cell types based on the location on the latent space.

We sample from the variational posterior using the reparametrization trick [30] as well as “mini-batches” from the dataset to compute unbiased estimate of the objective gradients’ with respect to the parameters. We use Adam [78] as a first-order stochastic optimizer to update the model parameters.

### Hyperparameters

For all harmonization tasks in this paper, we consistently use the same set of hyperparameters. Each network has exactly 2 fully-connected layers, with 128 nodes each. The number of latent dimensions is 10, the same as other algorithms for benchmarking purposes (e.g., the number of canonical correlation vectors used in Seurat Alignment). The activation functions between two hidden layers are all ReLU. We use a standard link function to parametrize the distribution parameters (exponential, logarithmic or softmax). Weights for the first hidden layer are shared between *f*_*w*_ and *f*_*h*_. We use Adam with *η* = 0.001 and *ϵ* = 0.01. We use deterministic warmup [79] and batch normalization [80] in order to learn an expressive model. When we train scANVI, we therefore assume that the data comes from a set of *C*_observed_ + *C*_unobserved_ populations, each generated by a different distribution of *z*_*n*_ values. This set includes the *C*_observed_ populations for which annotated cells are available, and *C*_unobserved_ population that accounts for cell types for which an annotation is not available to the algorithm. Ad-hoc training procedures for scANVI inference are described in **Supplementary Note 3**. In the case of a one single dataset, we use a 2 fully-connected layers with 256 hidden units classifier with *c* = 1 epochs of classifier training in between each variational update. In the case of transfer of labels, we use **Algorithm 2** with 1 fully-connected layers with 128 hidden units classifier, with *c* = 100 epochs of classifier training in between each variational update.

### Bayesian differential expression

#### scVI

For each gene *g* and pair of cells (*z*_*a*_, *z*_*b*_) with observed gene expression (*x*_*a*_, *x*_*b*_) and dataset identifier (*s*_*a*_, *s*_*b*_), we can formulate two mutually exclusive hypotheses:

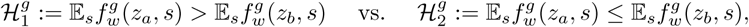

where the expectation 𝔼_*s*_ is taken with the empirical frequencies. Notably, we propose a hypothesis testing that do not calibrate the data to one batch but will find genes that are consistently differentially expressed. Evaluating which hypothesis is more probable amounts to evaluating a Bayes factor [81] (Bayesian generalization of the p-value) which is expressed as

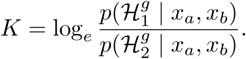

The sign of *K* indicates which of 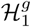 and 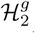 is more likely. Its magnitude is a significance level and throughout the paper, we consider a Bayes factor as strong evidence in favor of a hypothesis if |*K*| > 3 [82] (equivalent to an odds ratio of *exp*(3) ≈ 20). Notably, each of the probabilities in the likelihood ratio for *K* can be written as

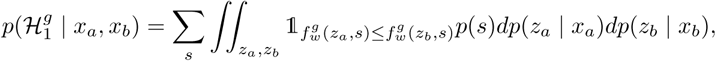

where *p*(*s*) designated the relative abundance of cells in batch *s* and all of the measures are lowdimensional. Since we cannot in principle achieve efficient posterior sampling, the naive Monte Carlo estimator obtained by replacing the real posterior *p*(*z* | *x*) by the variational posterior *qΦ*(*z* | *x*) is biased. The resulting Bayes factors are therefore approximate though yield very competitive performance, as explained in the original publication of scVI [29]. Since we assume that the cells are independently distributed, we can average the probabilities for the hypotheses across a large set of randomly sampled cell pairs, one from each subpopulation. The Bayes factor from the averaged probability will provide an estimate of whether cells from one subpopulation tend to express *g* at a higher frequency.

#### scANVI

In the case of scANVI, we need not rely on specific cells since labels are given during the training. We still use a generative model but with the following probability for 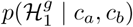 where *c*_*a*_ (resp. *c*_*b*_) is the first (resp. second) cell type of interest,

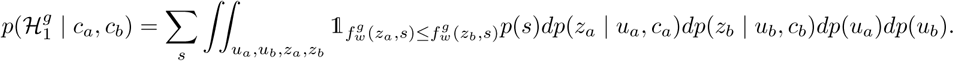

Notably, we draw here data from the prior distribution and not the posterior for given cells. As a consequence, these Bayes factors can be approximated in a unbiased fashion using a naive Monte Carlo estimator. We noticed in the case of the real dataset that the aggregate posterior on *u* might not perfectly match the prior for rare cell types. Consequently, we replaced the prior by the aggregate posterior for all the analysis in this manuscript.

#### Scalability

From a scalability perspective, our methods scales to a million cells in less than an our as reported in our original manuscript [29]. Remarkably, our inference procedure is scalable with respect to the number of batches (with number of floating point operations linear per dataset per iterations) while even the most efficiently implemented method such as Scanorama needs to compare each dataset pairs (quadratic). Our runtime on the large dataset from Scanorama is 42 min for 250 iterations over 100K cells from 26 datasets, still 30 min longer than their algorithm. We expect that further work on implementation details such as GPU usage efficiency, memory loading, data format and iteration monitoring will help close the computational gap between these algorithms.

### Datasets

We report an extensive list of datasets at Supplementary Table 1. For all UMI based datasets we took the raw counts without any normalization as input to scVI.

#### Gene Selection

A common practice in data harmonization is to perform gene selection prior to harmonization. This assumption is critical when the number of genes that can be taken into account by the algorithm is small and potentially biological signal could be lost. scVI is however designed for large datasets which do not fall into the high-dimensional statistics data regime [29]. Remarkably, there is no need for crude gene filtering as part of our pipeline and we adopt it as part of this publication only for concerns of fairness in benchmarking. For real datasets, we calculated the dispersion (variance to mean ratio) for all genes using Seurat in each dataset and selected *g* = 1, 000 genes with the highest dispersion from each. The performance of scVI is not as affected by gene set and we use the same gene selection scheme as in [13] to ensure fairness in our comparison. We then took the union of these gene list as input to Seurat Alignment, MNN and scANVI. One exception is the differential expression study for which we kept the gene set (*g* = 3, 346) to have it match the bulk reference as in [29].

#### Cell type labeling for the Tabula Muris Dataset

For the Tabula Muris dataset, cell types are defined by first reducing the dimensions of the data by principal component analysis and then performing nearest-neighbor-graph-based clustering. The labels for Smart-Seq2 and 10x data are derived independently. All cells in both dataset are labeled, but there is also a possibility that they are mislabelled since the labels are computationally derived. Since cells used in Smart-Seq2 are first FACS sorted into each plate, some cell types might have been lost during the sorting process, resulting in incomplete overlap in cell types between the two datasets.

#### Hierarchical cell type labeling for the mouse nervous system dataset

The multilevel labels are generated through an iterative process that is described in detail in the original publication [60]. The clustering was performed with strict quality filters, takes into account anatomical information and were validated at different levels using existing scRNAseq dataset, osmFISH, RNAscope and others. The cell types taxonomy is derived differently for each level and the details can be found in the original publication. Cell type clusters were obtained by Louvain clustering on a multiscale *k*NN graph and DBSCAN. The first level separates neurons and non-neuronal cells. The second level separates peripheral neuronal system from central neuronal system. The third layer separate anterior posterior domain, and the fourth layer is split by excitatory versus inhibitory neurotransmitter. At this level, all cells are divided into 39 subsets, each corresponding to a coarser cell type definition. Then, within each subset the authors defined N=28 enriched genes and used linkage (correlation distance and Ward method) to construct the dendrogram.

#### Normalization of SmartSeq2 data

For the MarrowMT-ss2 dataset, we normalized the read counts per gene by relative transcript length (average transcript lengths of a gene divided by average gene length over all genes), and subsequently took the integer part of the normalized count. This is different from standard normalization procedures in that we do not normalize by cell size because cell size normalization can be performed by scVI. And we only keep the integer part of the counts, due to the distributional assumptions made by scVI. The scVI model can to be extended to fit data with amplification bias, however we have not done so for this paper and thus have to perform this normalization heuristic.

#### Normalization of CITE-seq data

Since we did not explicitly model the CITE-seq data in our models, we normalized it by fitting a Gaussian mixture model to each individual protein with two components. We then transformed each individual protein count as 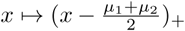 where *µ*_1_ and *µ*_2_ designate the mean of the mixtures and. _+_ is the positive part of a real number.

#### Simulation using SymSim

First we simulated the true expression matrix for 20,000 cells from 5 cell types using the function SimulateTrueCounts. We then randomly split the cells into two batches. We then added noise to the data the function True2ObservedCounts with the parameters

~~~
Batch 1: protocol=“UMI”, alpha_mean=0.03, alpha_sd=0.009, gene_len=gene_len, depth_mean=5e5, depth_sd=1.5e4
Batch 2: protocol=“UMI”, alpha_mean=0.1, alpha_sd=0.03, gene_len=gene_len, depth_mean=1e6, depth_sd=1.5e5
~~~

#### Continuous Simulation using SymSim

First we simulated the true expression matrix for a tree with 5 cell types using the function SimulateTrueCounts. Instead of sampling cells only from the leaf populations, we uniformly sample cells along all branches by using the parameter evf type=“continuous”. We then added noise to the data with the function True2ObservedCounts with the parameters

~~~
protocol=“nonUMI”, alpha_mean = 0.1, alpha_sd = 0.05, rate_2PCR = 0.7, nPCR1 = 16,depth_mean=1e5, depth_sd=3e3
~~~

### Algorithms for benchmarking

#### Seurat Alignment

We applied the Seurat Alignment procedure from the R package Seurat V2.3.3. The number of canonical correlation vectors is 10 for all the datasets, which is also identical to the number of latent dimensions used for scVI and scANVI.

#### Seurat PCA

We applied the Seurat PCA procedure from the R package Seurat V2.3.3. This method is a simple PCA based after normalization by Seurat. Seurat PCA is used to obtain the individual dataset latent space to evaluate the *k*-nearest neighbors purity for all non-scVI based methods. The number of principal components is 10.

#### Matching Mutual Nearest Neighbors

We used the mnnCorrect function from https://rdrr.io/bioc/scran/man/mnnCorrect.html with default parameters. In order to compare with other methods, we applied a PCA with 10 principal components on the output of the batch-corrected gene expression matrix.

#### scmap

We applied the scmap-cluster procedure from the R package scmap. As the scmap manuscript insists heavily on why the M3Drop [83] gene filtering procedure is crucial to overcome batch effects and yield accurate mapping, we let scmap choose its default number of genes (*g* = 500) with this method.

#### ComBat

We used the R package sva with default parameters.

#### UMAP

We used the umap class from the UMAP package with a default parameters and spread=2.

#### DBSCAN

We used the DBSCAN algorithm from the Python package from the python package scikit-learn V0.19.1 and we searched for an optimal hyperparameter combination by a grid search over `eps` and `min_samples` from the range of 0.1 − 2 and 5 − 100 respectively. Although some combinations of parameters yield more than one clusters, the smaller clusters comprise of less than 1% of the data. We then evaluated DBSCAN with `eps=1.23, min\_samples=10` and default values for all other hyper-parameters.

#### PhenoGraph

We used the phenograph.cluster function from the Python package PhenoGraph 1.5.2 downloaded from https://github.com/jacoblevine/PhenoGraph with its default parameters.

#### CORAL

We used the implementation from https://github.com/jindongwang/transferlearning/tree/master/code/traditional/CORAL.

#### MAGAN

We used the implementation from https://github.com/KrishnaswamyLab/MAGAN

### Evaluations metrics

#### Entropy of batch mixing

Fix a similarity matrix for the cells and take *U* to be a uniform random variable on the population of cells. Take *B*_*U*_ the empirical frequencies for the 50 nearest neighbors of cell *U* being a in batch *b*. Report the entropy of this categorical variable and average over *T* = 100 values of *U*.

#### *k*-nearest neighbors purity

Compute two similarity matrices for cells from the first batch, one from the latent space obtained with only cells from the first batch and the other from the latent space obtained using both batches of cells. We always rely on the euclidean distance on the latent space. Take the average ratio of the intersection of the *k*-nearest neighbors graph from each similarity matrix over their union. Compute the same statistic for cells from the other batch and report the average of the two.

#### Weighted and unweighted accuracy

We can evaluate accuracy of cell type classification by comparing prediction to the previously published labels. The unweighted accuracy is the percentage of cells that have the correct label. The weighted accuracy is when the accuracy is calculated for each cell type first and averaged across cell types. The weighted accuracy assigns the same weight to each cell type and thus weighs correct prediction of rare cell types more heavily than the unweighted accuracy. All accuracies reported in this paper is the weighted accuracy.

#### Maximum Posterior Probability

We evaluate the performance of the scANVI classifier at transferring labels from an annotated dataset to an unannotated dataset by looking at the maximum posterior probability for the observed classes. By default scANVI classifier sets the number of classes to the same number of cell types in the merged dataset. In the case of *N* observed labels from the annotated dataset and one unannotated dataset (thus the cell type label is “Unlabeled”) scANVI assumes *N* + 1 classes. For each cell, scANVI assigns a posterior probability for each of the *N* + 1 classes. The maximum posterior probability for the observed classes is the highest probability of a cell being assigned to one of the *N* observed classes.

### Signature for sub-division of T cells in human PBMCs

#### Gene sets

For ranking the cells, we used both positive and negative sets of genes:

- **CD4 Regulatory:** *GITR*+ *CTLA4* + *FOXP3* + *CD25* + *S100A4* - *CD45* - *CD8B* -
- **CD4 Naive:** *CCR7* + *CD4* + *S100A4* - *CD45* - *FOXP3* - *IL2RA*- *CD69* -
- **CD4 Memory:** *S100A4* + *CD25* - *FOXP3* - *GITR*- *CCR7* -
- **CD8 Naive:** *CD8B* + *CCR7* + *CD4* -

#### Signature calculus

To compute the signature of a cell, we followed the normalization procedure from [24] which consists in dividing by total numbers of UMIs, applying a entry-wise transformation *x* ↦ log(1 + 10^4^*x*) and *z*-score normalization for each gene. Then, we aggregated over the genes of interest for each cell by applying the sign from the gene-set and averaging.

### Software Availability

An open-source software implementation of scVI is available on Github (https://github.com/YosefLab/scVI). All code for reproducing results and figures in this manuscript is deposited at https://doi.org/10.5281/zenodo.2529945.

### Data availability

All of the datasets analyzed in this manuscript are public and referenced at https://github.com/chenlingantelope/HarmonizationSCANVI.

---

**Figure 1:**
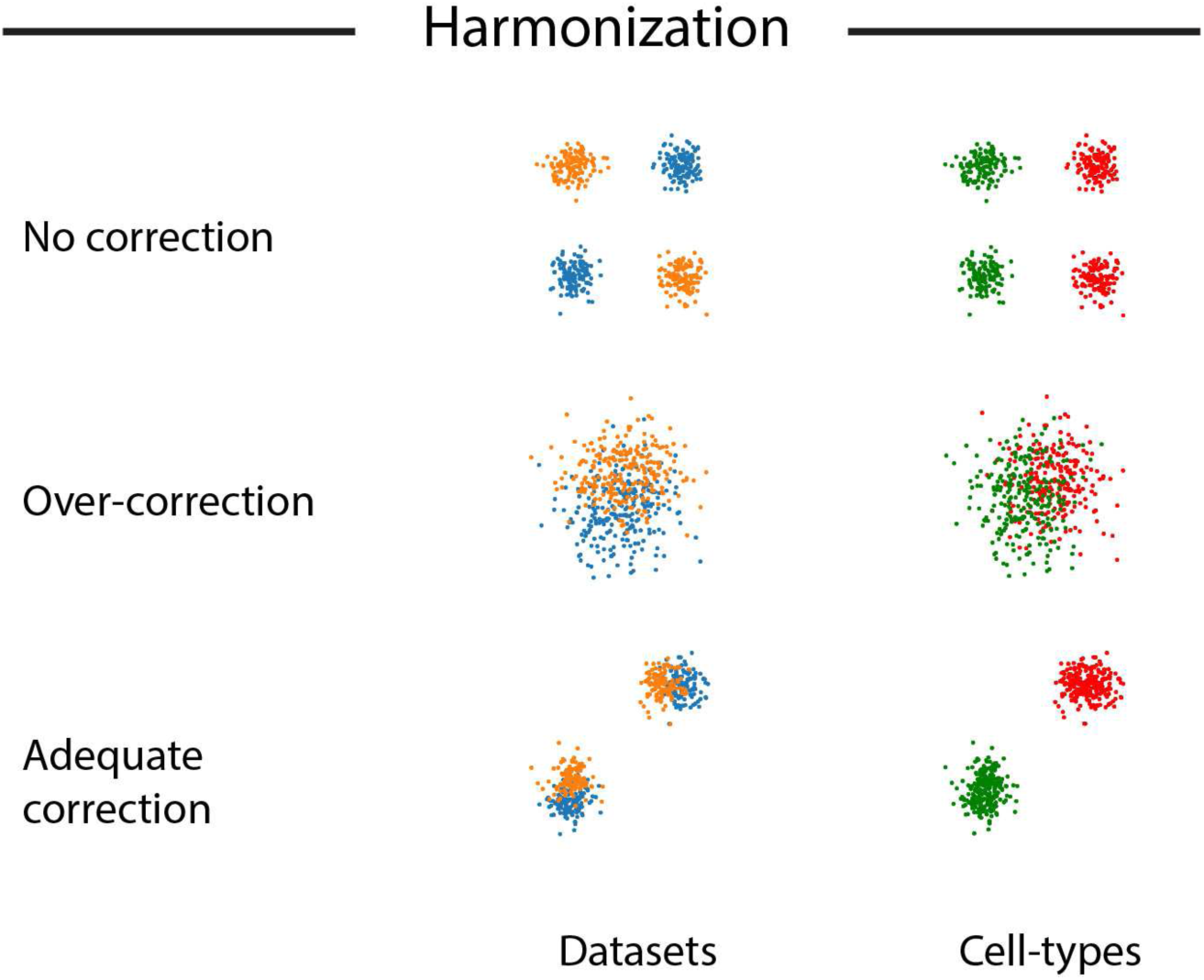
Schematic of the data harmonization problem. We are provided with two datasets (orange and blue), each consisting of two cell types (red and green). Our evaluation for the harmonization problem consists of two objectives: (1) mixing the two datasets well and (2) retaining the original structure in each dataset. Scenario 1 (top) is the case of under correction where objective (2) is achieved while objective (1) is not. Scenario 2 (middle) is the case of over correction where objective (1) is improved while objective (2) becomes worse. The bottom panel shows the desired scenario of mixing the datasets well while retaining the biological signal.

**Figure 2:**
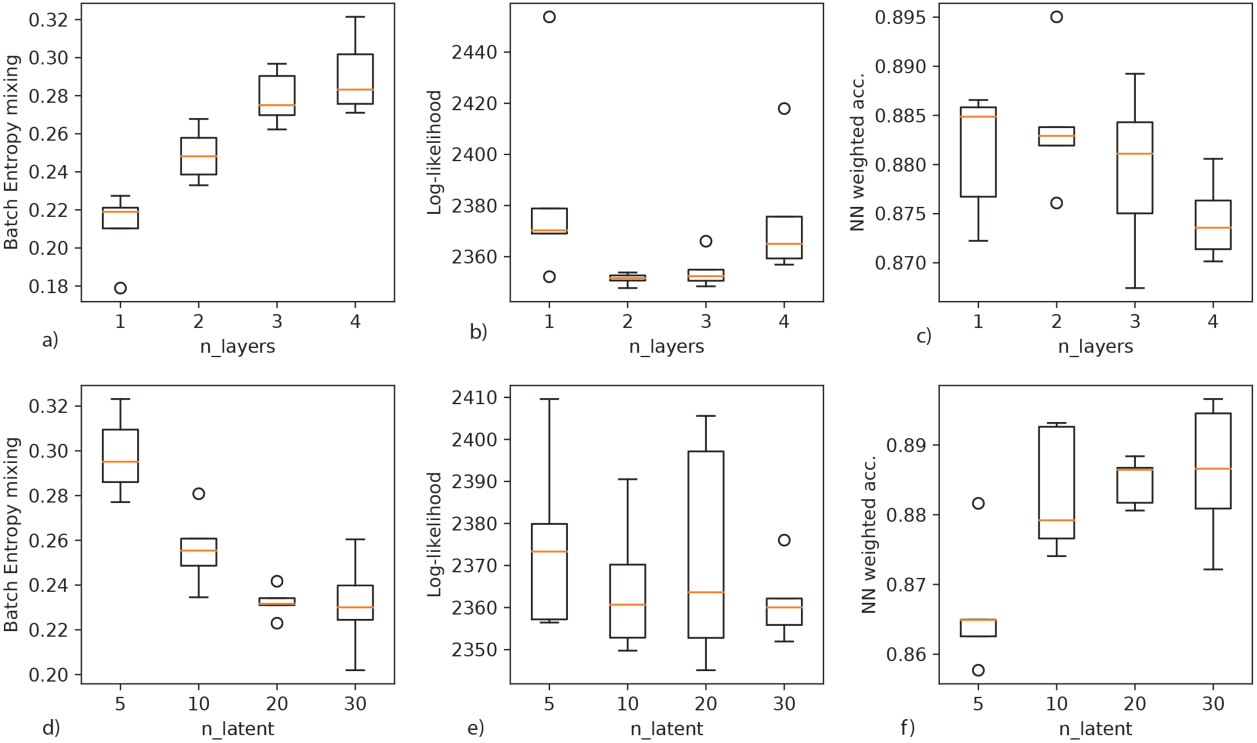
Robustness analysis for harmonization of the pair of datasets MarrowMT-10x / MarrowMT-ss2 with scVI. (*a* − *c*) We augment the number of hidden layers in the neural network *f*_*w*_ and track across *n* = 5 random initializations for the batch entropy mixing (*a*), the held-out log likelihood (*b*) and the weighted accuracy of a nearest neighbor classifier on the latent space (*c*). (*d* − *f*) We increase the number of dimensions for the latent variable *z* and track across *n* = 5 random initialization the batch entropy mixing (*d*), the held-out log likelihood (*e*) and the weighted accuracy of a nearest neighbor classifier on the latent space (*f*)

**Figure 3:**
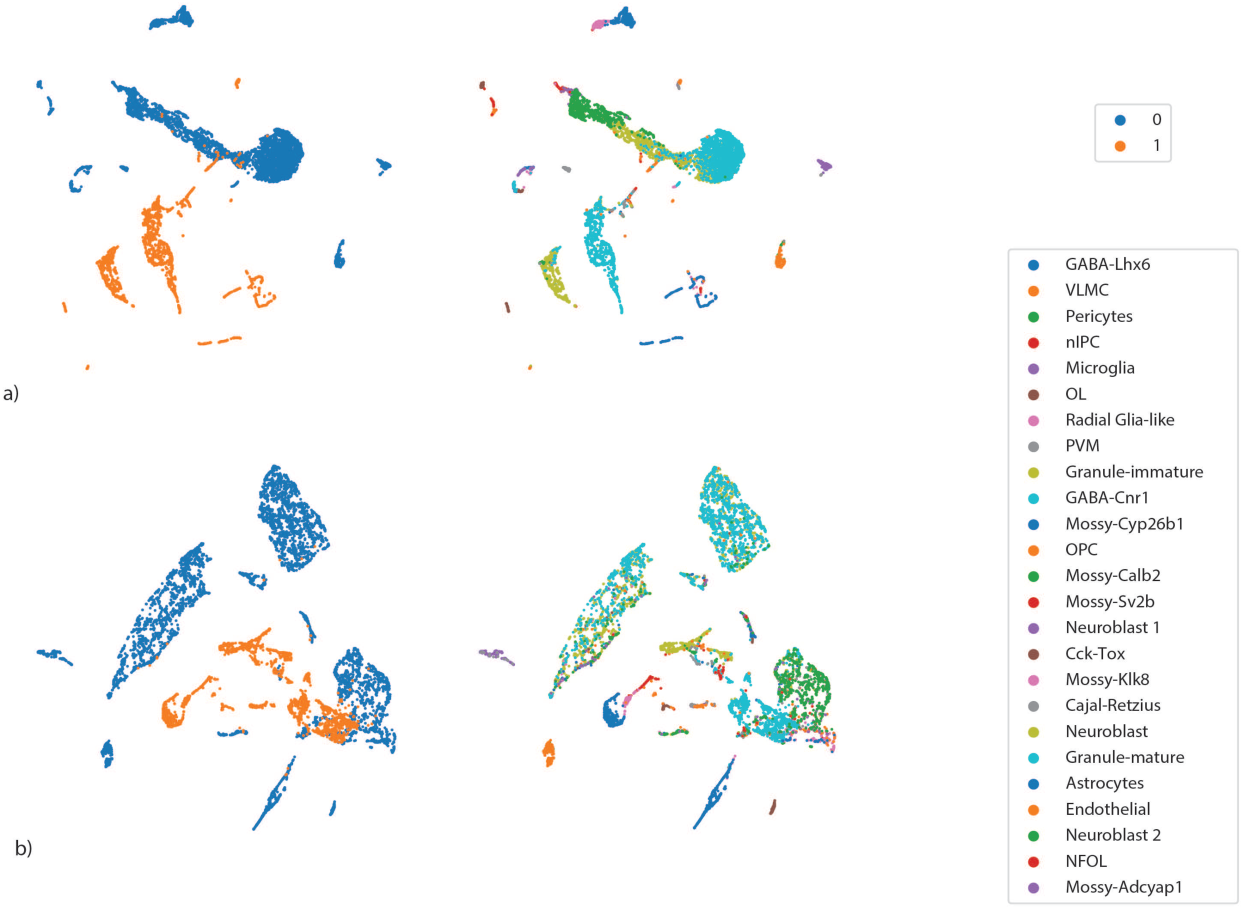
Results of single-cell harmonization with MAGAN on the DentateGyrus pair of datasets. Using MAGAN, we projected the first dataset into the second one (*a*) and vice-versa (*b*).

**Figure 4:**
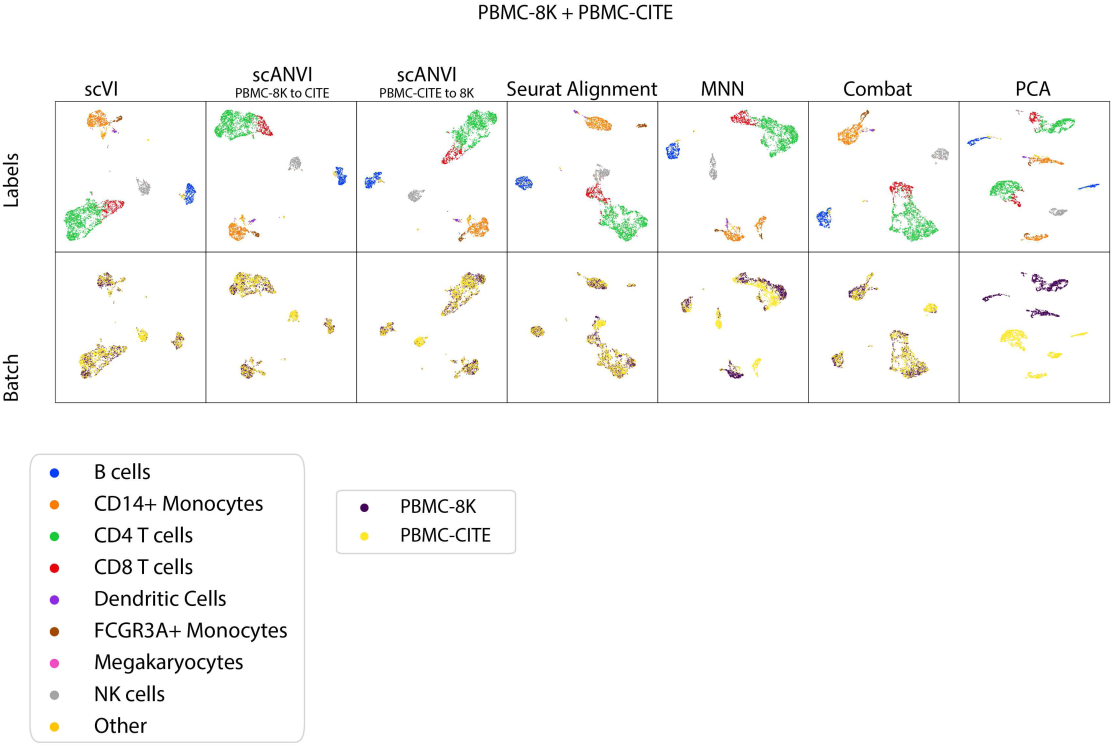
Visualization of the benchmark PBMC-8K / PBMC-CITE. all positions for the scatter plots are derived using UMAP on the latent space of interest.

**Figure 5:**
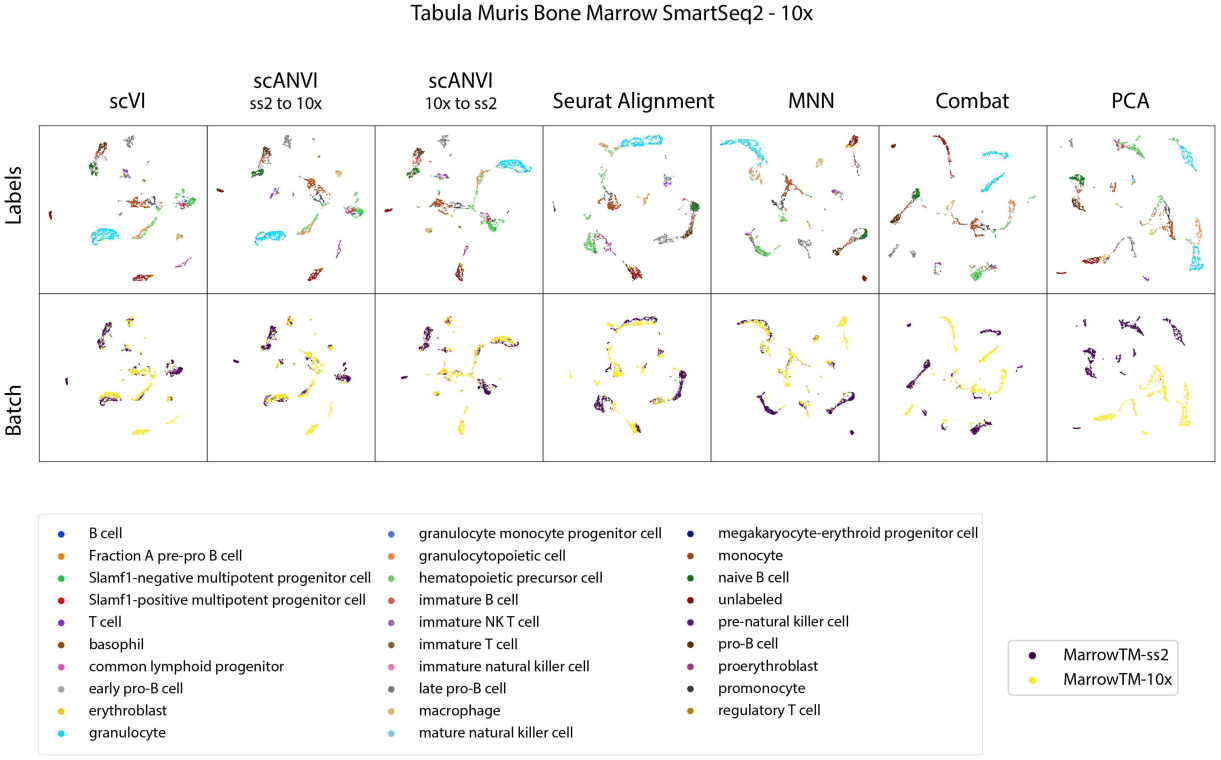
Visualization of the benchmark MarrowMT 10x / ss2. all positions for the scatter plots are derived using UMAP on the latent space of interest.

**Figure 6:**
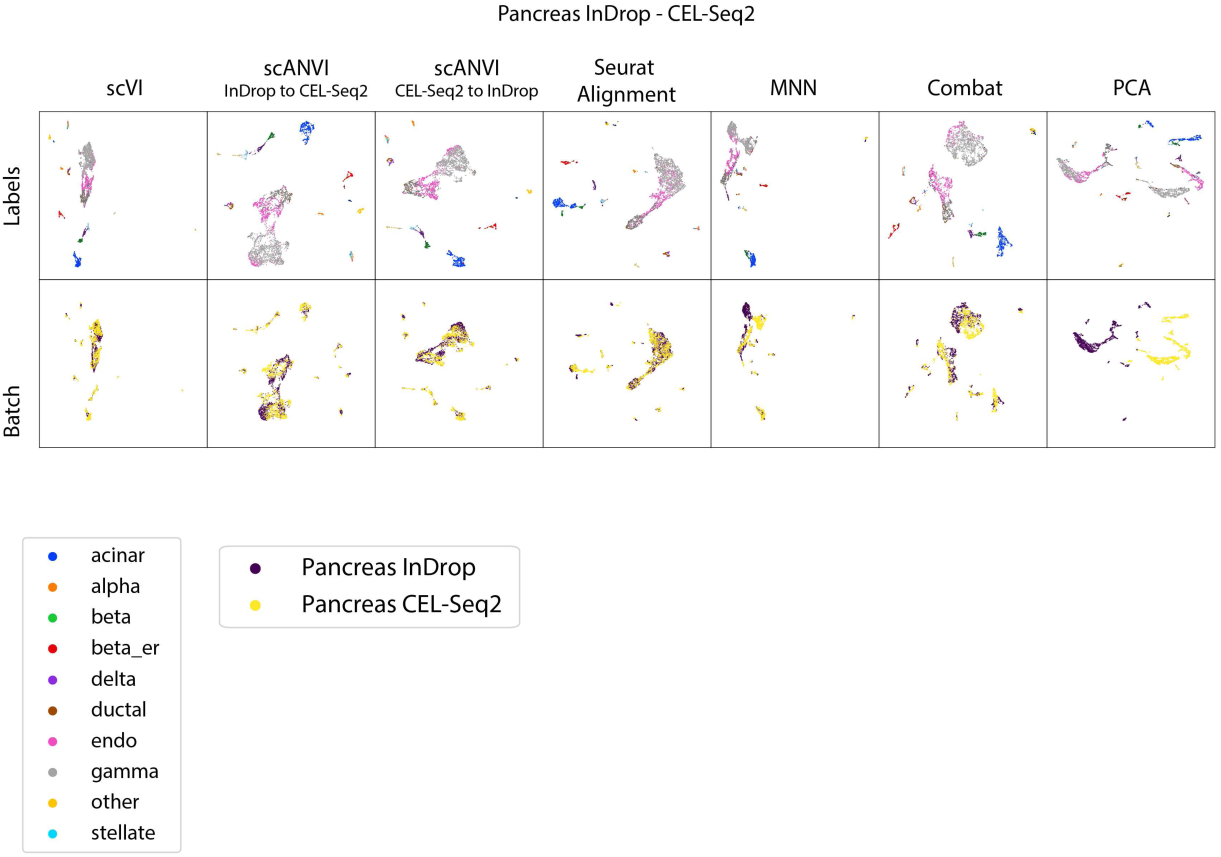
Visualization of the benchmark Pancreas InDrop / CEL-Seq2. all positions for the scatter plots are derived using UMAP on the latent space of interest.

**Figure 7:**
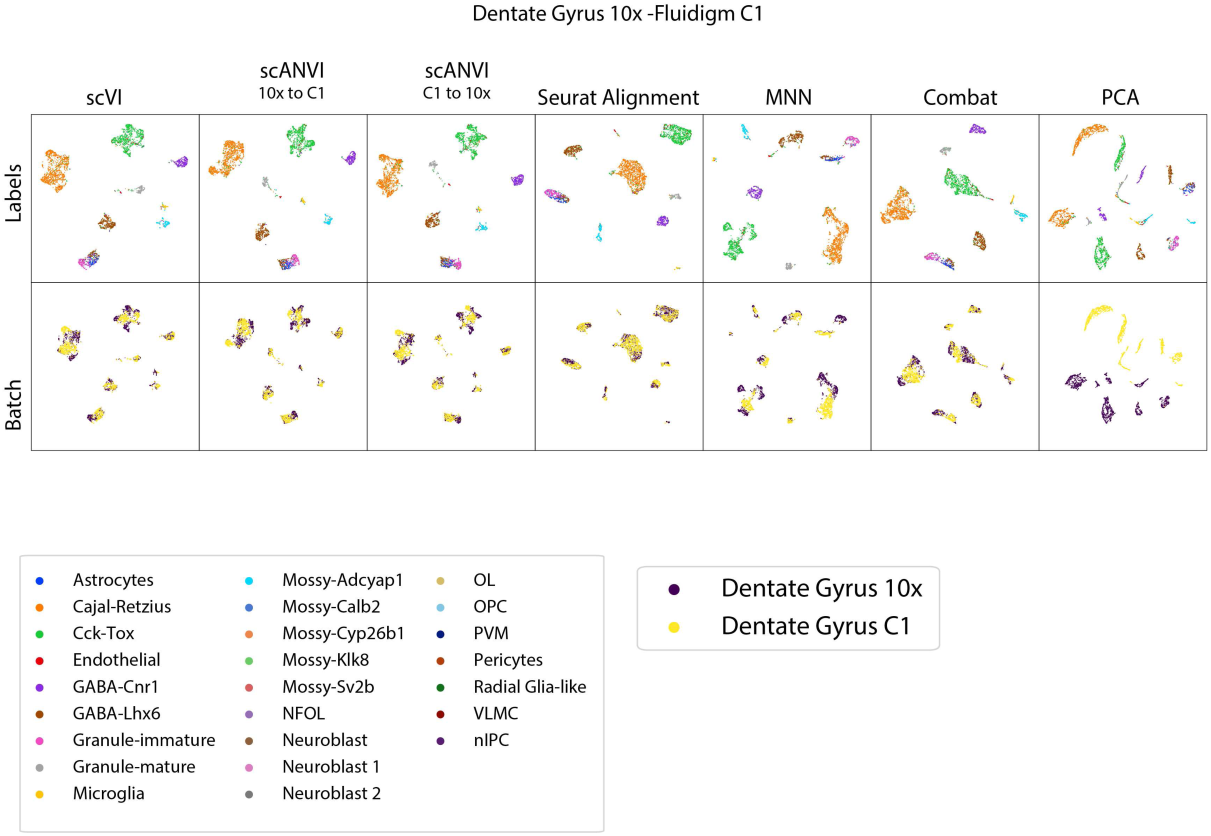
Visualization of the benchmark Dentate Gyrus 10X - Fluidigm C1. all positions for the scatter plots are derived using UMAP on the latent space of interest.

**Figure 8:**
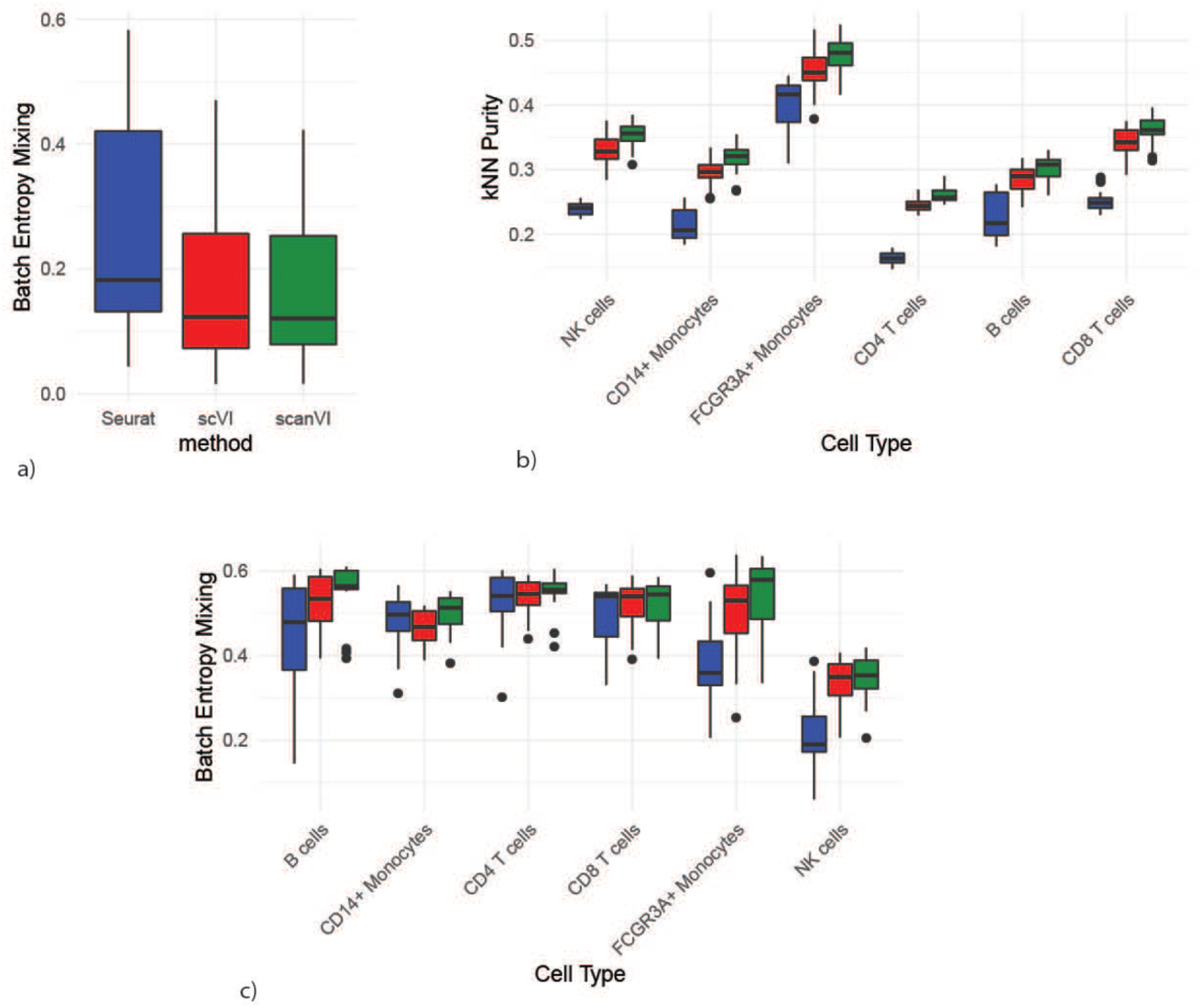
Supplementary study of harmonizing datasets with different cellular composition. We show here the case where each of the two datasets has a unique cell types and share all the others. For each box plot, we report over all the possible combinations of left-out cell types. (a) Batch entropy mixing on the unique populations (lower is better). (b) Conservation of the original structure across all populations (unique and non-unique; higher is better). (c) Batch Entropy mixing on the non-unique populations (higher is better). The boxplots are standard Tukey boxplots where the box is delineated by the first and third quartile and the whisker lines are the first and third quartile plus minus 1.5 times the box height. The dots are outliers that fall above or below the whisker lines.

**Figure 9:**
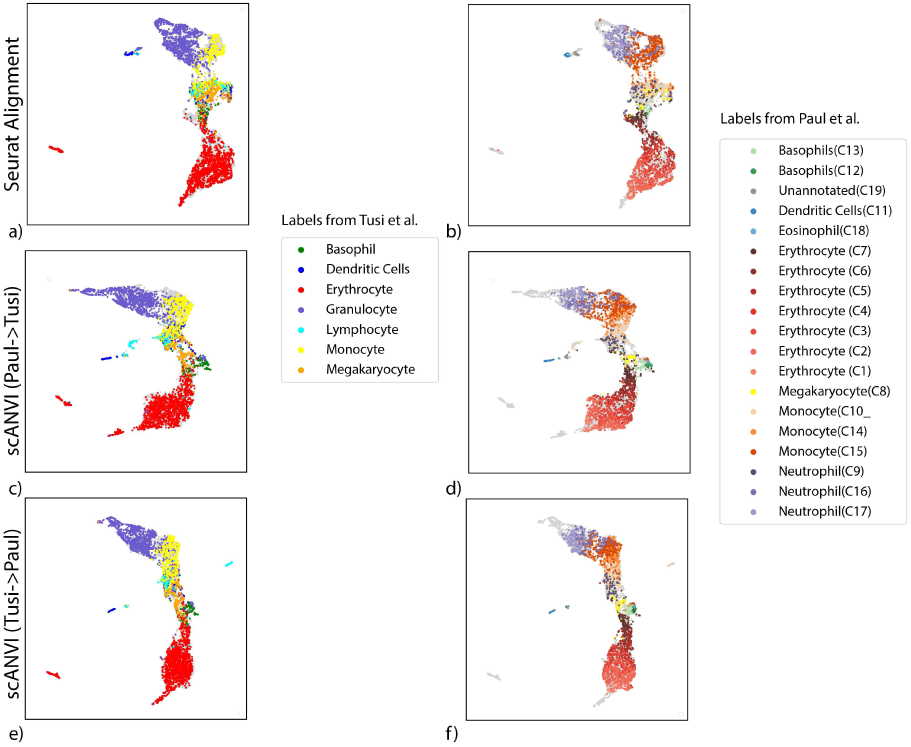
(*a* − *b*) Continuous trajectory obtained by the Seurat Alignment procedure for the HEMATO-Tusi and the HEMATO-Paul datasets. (*c* − *d*) Continuous trajectory obtained by the scANVI using the Tusi cell type labels for semi-supervision. (*e* − *f*) Continuous trajectory obtained by the scANVI using the Paul cluster labels for semi-supervision. All locations for scatter plots are computed via UMAP in their respective latent space.

**Figure 10:**
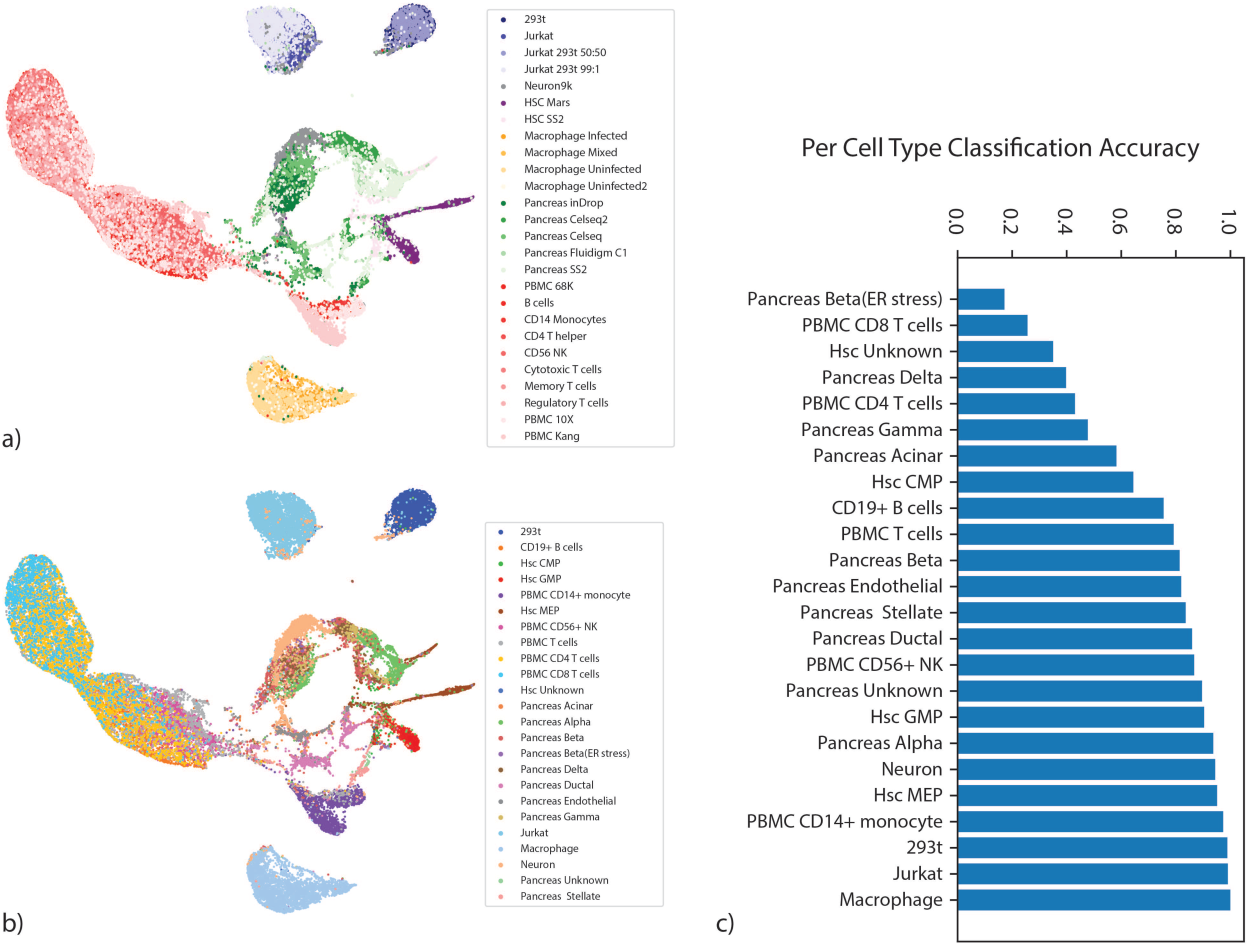
Large Scale data integration with scVI. (*a* − *b*) UMAP visualization of the scVI latent space colored by datasets (*a*) and by cell types (*b*). (*c*) accuracy of a nearest neighbor classifier based on scVI latent space

**Figure 11:**
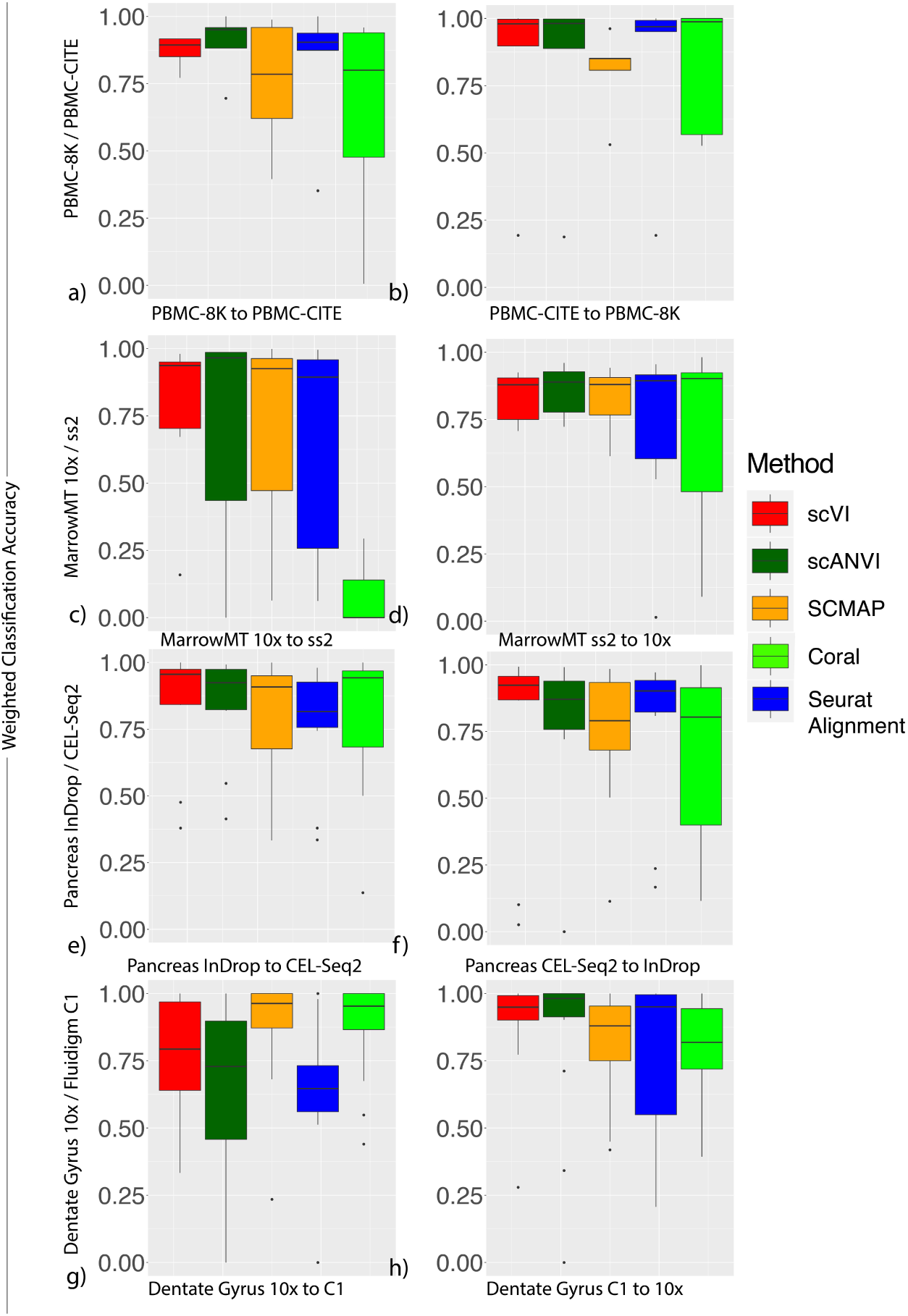
Annotation results for the PBMC-8K / PBMC-CITE (*a* − *b*), the MarrowMT-10X / MarrowMT-SS2 datasets datasets (*c* − *d*), Pancreas InDrop-CITESeq (*e* − *f*) and Dentate Gyrus 10X / Fluidigm C1 (*g* − *h*). Accuracies for transferring annotations from one dataset to another from a *k*-nearest neighbor classifier on Seurat Alignment, *k*-nearest neighbor classifier on scVI latent space, scANVI, SCMAP and CORAL classifier are shown. The aggregated results across for cell types that are shared between the two datasets is shown in box plots.

**Figure 12:**
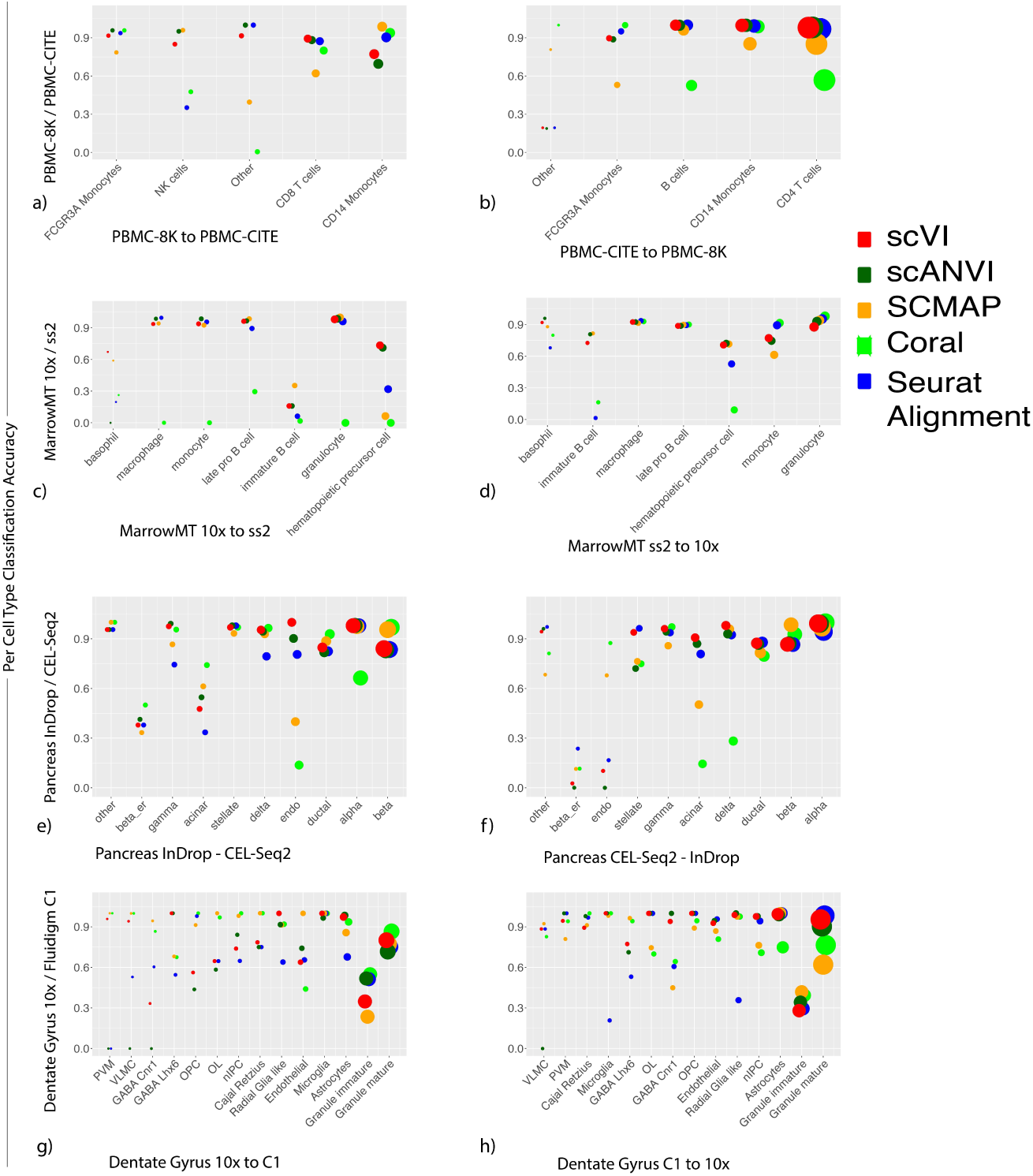
Annotation results for the PBMC-8K / PBMC-CITE (*a* − *b*), the MarrowMT-10X / MarrowMT-SS2 datasets datasets (*c* − *d*), Pancreas InDrop-CITESeq (*e* − *f*) and Dentate Gyrus 10X / Fluidigm C1 (*g* − *h*). Accuracies for transferring annotations from one dataset to another from a *k*-nearest neighbors classifier on Seurat Alignment, *k*-nearest neighbors classifier on scVI latent space, scANVI, SCMAP and CORAL classifier are shown. The prediction accuracy for each cell type that is shared between the two datasets is shown on the y-axis and the size of the dots are proportional to the proportion of a cell type in the total population.

**Figure 13:**
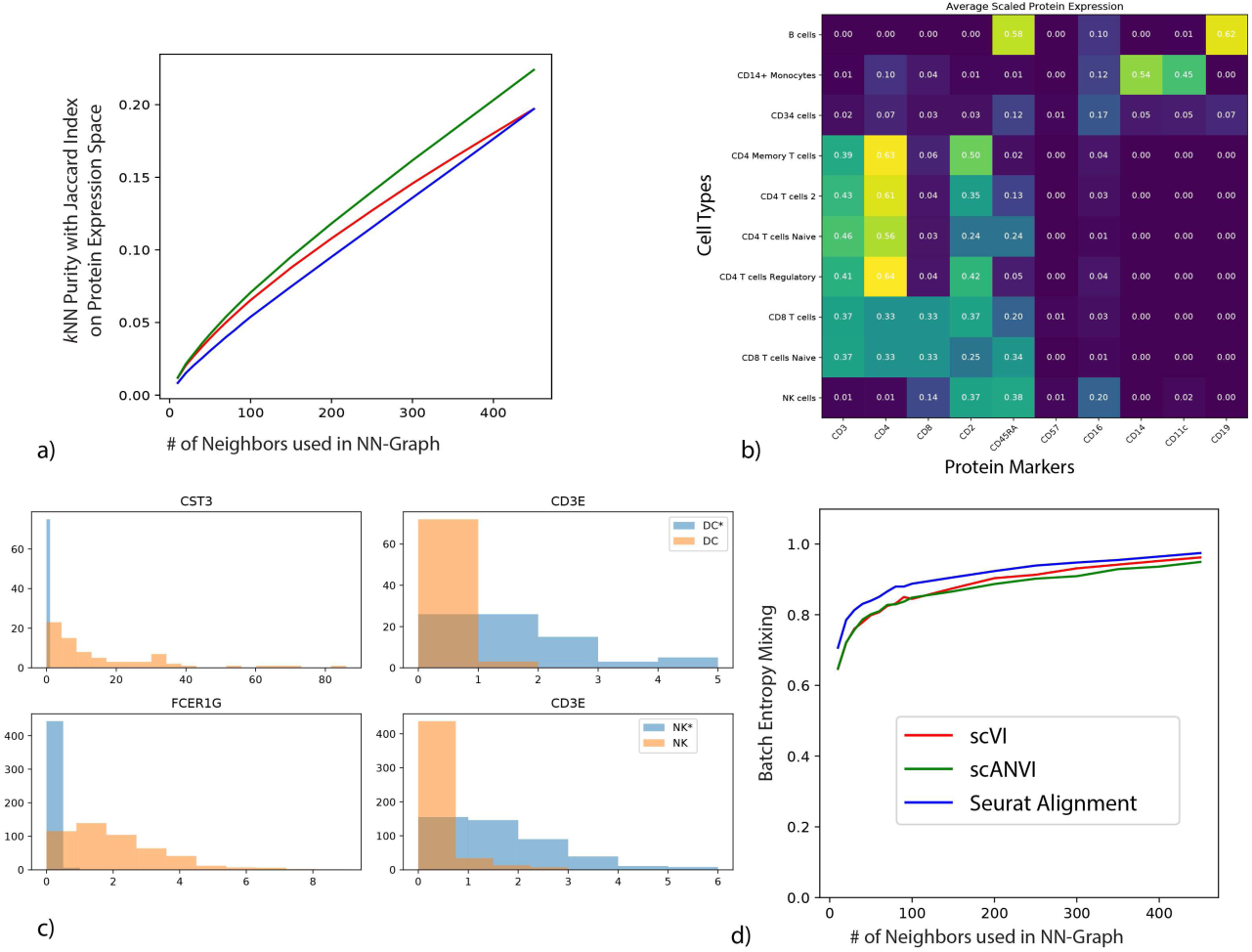
(*a*) *k*-nearest neighbors purity of the merged latent space on the protein expression space as a function of the size of the neighborhood. (*b*) Protein expression heatmap showing consistency of PBMC-sorted labels and protein expression in PBMC-CITE. The protein expression per cell type is based on *k*-nearest neighbors imputation from the harmonized latent space obtained from scANVI trained with pure population labels. (*c*) We select individual cells that were labeled as Dendritic cells or Natural Killer cells in the original publication of the respective datasets, and compare the raw transcript count from cells inside the scANVI T cells cluster (DC*, NK*) against cells outside the T cells cluster (DC, NK). The expression of marker genes suggest that DC* and NK* is more likely to be T cells and thus the scANVI latent space is more accurate. (*d*) The batch entropy mixing of the three datasets in scVI, scANVI and Seurat Alignment merged space.

**Figure 14:**
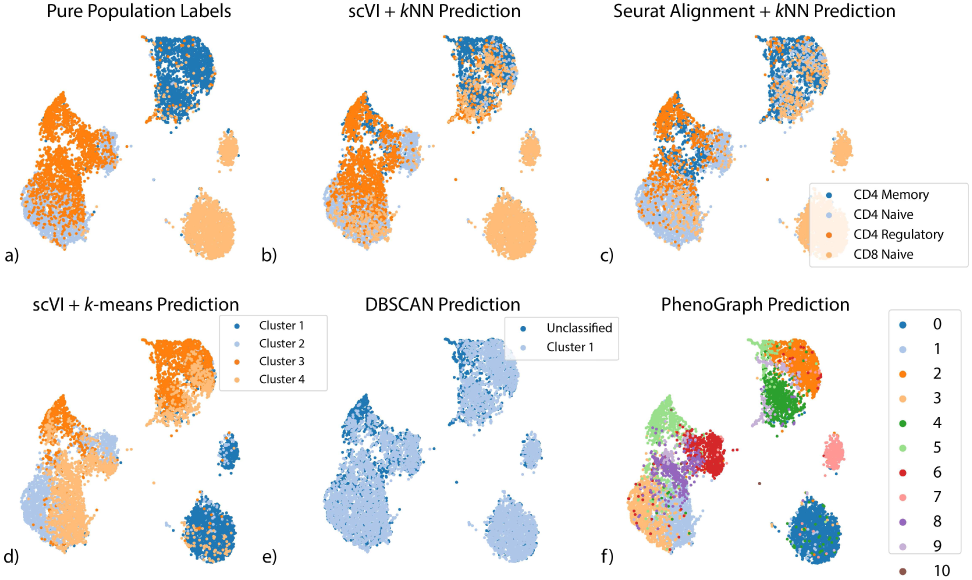
Other methods of classifying T-cell subsets of the PBMC-Pure dataset. Coordinates for the scatter plots are derived from UMAP embedding based on the latent space of scANVI. (*a*) Ground truth labels from the purified PBMC populations (*b*) *k*-nearest neighbors classification labels when applied on scVI latent space from the seed set of cells (*c*) *k*-nearest neighbors classification labels when applied on Seurat Alignment latent space (*d*) *k*-means clustering based labels when applied to scVI latent space (*e*) DBSCAN clustering based labels when applied to scVI latent space. DBSCAN returns only one cluster but return some cells as unclassified. (*f*) PhenoGraph clusters on scVI latent space

**Figure 15:**
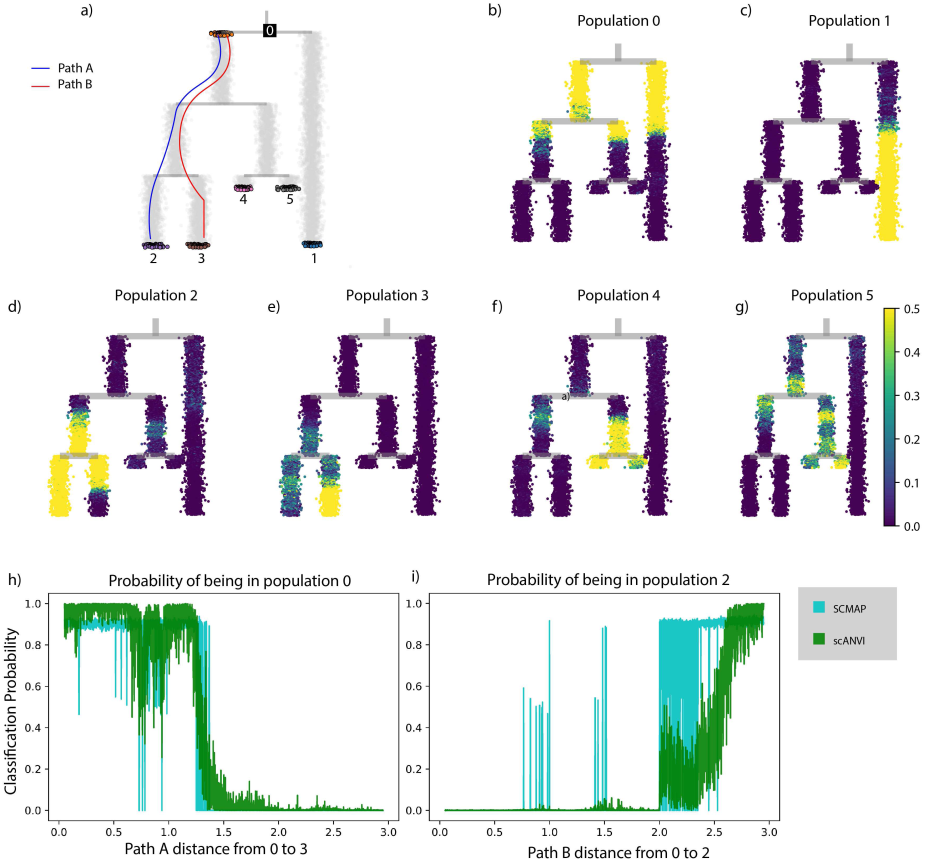
Continuous trajectory simulated using SymSim. (*a*) Tree structure from which the cells are sampled. Each grey dot represent a cell sampled along the trajectory. Colored dots with a black edge are treated as labeled, while the others are treated as unlabeled. Each path simulates a continuous phenotypical variation. (*b* − *g*) The same tree with each cell colored by the posterior probability of being assigned to a specific label. (*h* − *i*) Another visualization of the gradual change of posterior probability by plotting the posterior probability of root (*h*) and population 3 (*i*). The x-axis represents the pathwise distance (paths are defined in (*a*)), and the y-axis represents the probability, or confidence of the assignment.

**Figure 16:**
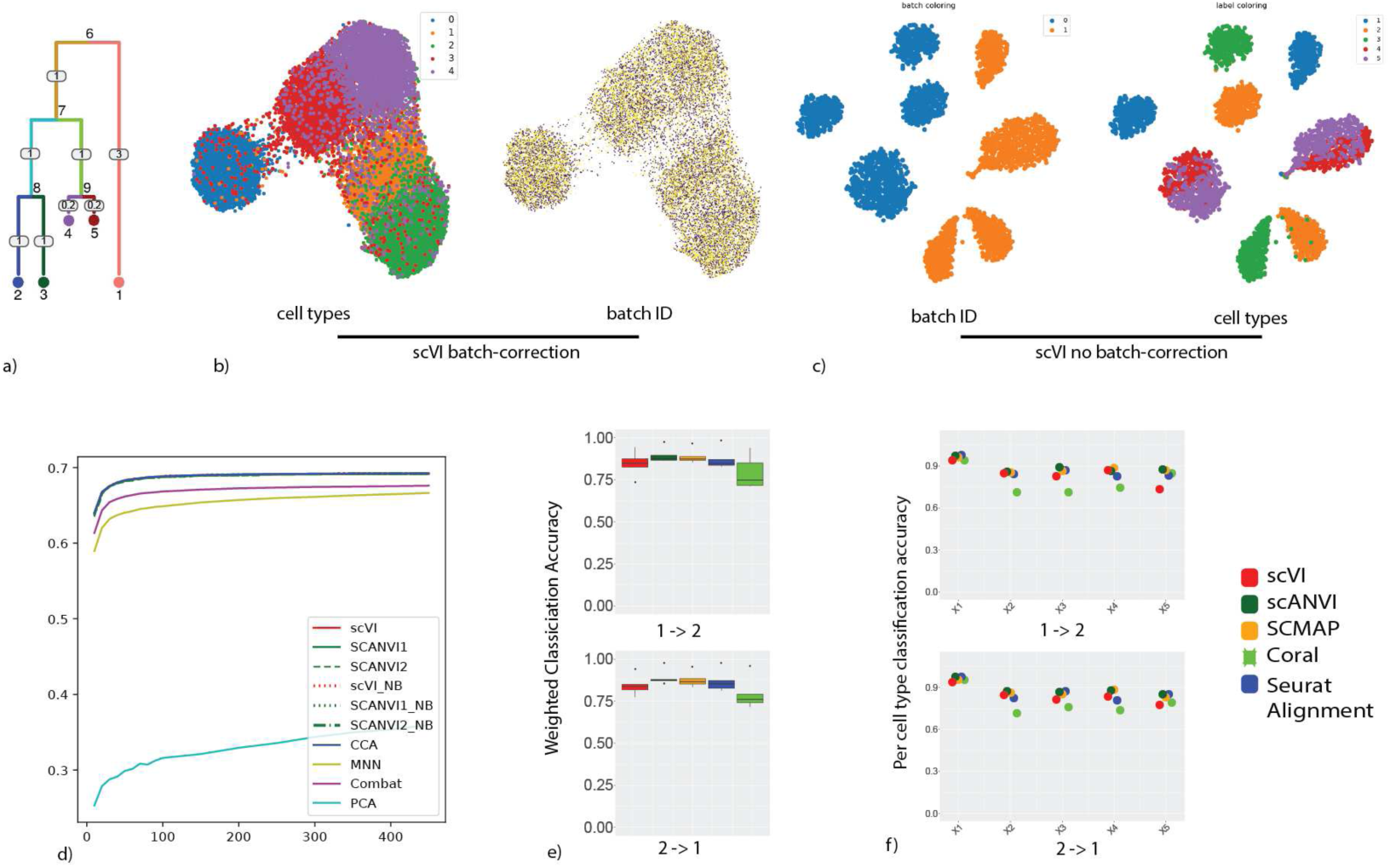
Presentation of the simulated dataset used for differential expression benchmarking. (a) The tree used to sample the cells in SymSim. We sample cells from the five leaves nodes representing five different cell types derived from the same root node. (b) UMAP of scVI latent space colored by cell types and batch identifier (c) UMAP of scVI without batch correction, proving that the data is indeed subject to batch effects. (d) Batch entropy mixing for all the algorithms (e) Weighted classification accuracy using a nearest-neighbor method on the latent space (f) Per cell type classification accuracy for the label transfer.

**Figure 17:**
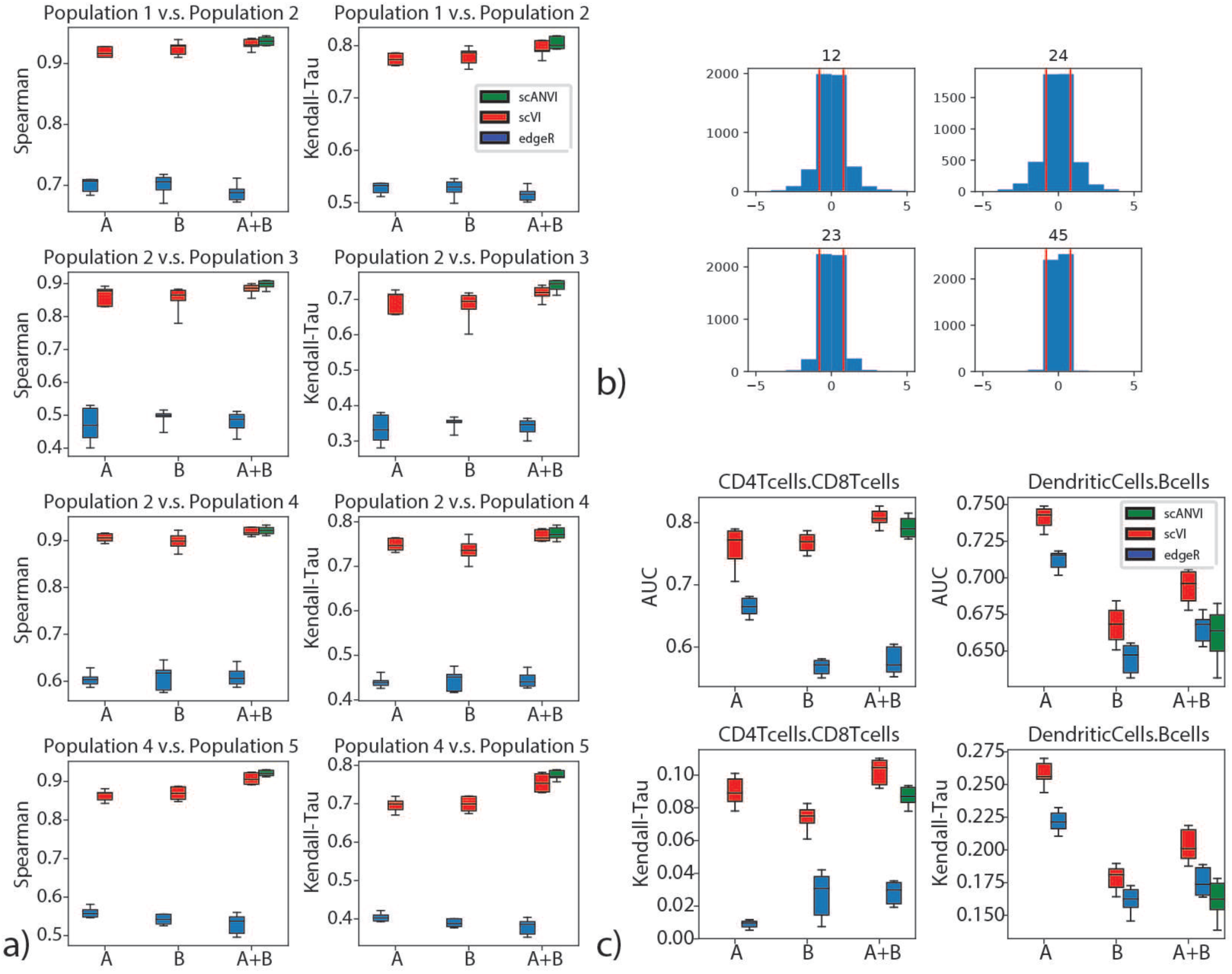
Differential Expression on multiple datasets with scVI. (*a*) Evaluation of consistency with rank correlation is shown for comparisons of multiple pairs of cell types in the simulated data. The pairs of cells are chosen to represent different levels of distance on the tree as in Figure 16a. The pairs of population from most distant to least distant are ‘12’, ‘24’, ‘23’, ‘45’. (*b*) distribution of true log fold change between all pairs of cell types for the simulated data. (*c*) Evaluation of consistency with the AUROC and Kendal Tau metric is shown for comparisons of CD4 vs CD8 T cells and B cells vs Dendritic cells on the PBMC-8K only (A), the PBMC-68k only (B) and the merged PBMC-8K / PBMC-68K (A+B) for scVI and edgeR. Error bars are obtained by multiple subsampling of the data to show robustness. boxplots are standard Tukey boxplots where the box is delineated by the first and third quartile and the whisker lines are the first and third quartile plus minus 1.5 times the box height. The dots are outliers that fall above or below the whisker lines.

**Figure 18:**
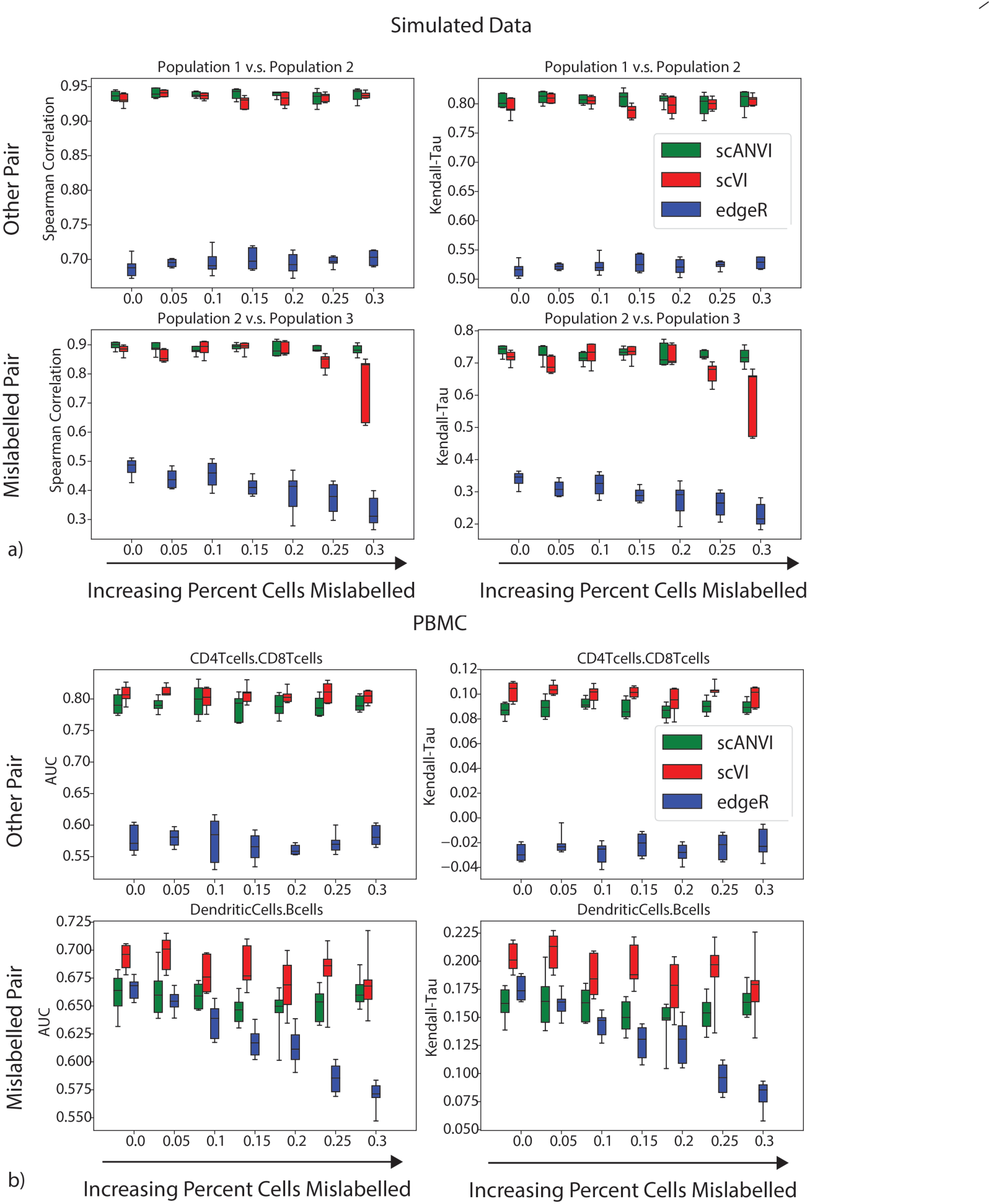
Mislabeling experiment in differential expression with (*a*) PBMC8K and PBMC68K datasets and (*b*) SymSim simulated datasets. Within each panel: Top: differential expression results for the correctly labeled population pair (CD4 T cells - CD8 T cells in (*a*) and 1-2 in (*b*)). Bottom: differential expression results for the mislabelled population pair (Dendritic Cells and B cells in (*a*) and 2-3 in (*b*)). For all, x-axis represents the proportion of flipped labels.

**Figure 19:**
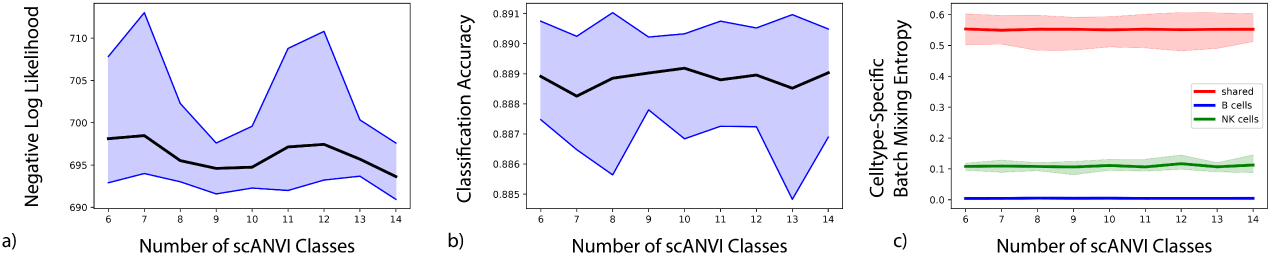
The effect of the choice of number of classes on the scANVI model likelihood (*a*), classification accuracy (*b*) and batch entropy mixing (*c*). We trained scANVI using PBMC8K as the labelled dataset, and varied the number of classes in scANVI from 6(true number of labelled cell types) to 14. The thicker line show the mean of 9 replicates, while the colored shading show the 95% confidence interval. We used a subsampled PBMC8KCITE dataset, where NK cells are removed from the PBMC8K dataset and B cells are removed from the PBMC-CITE dataset. As we expect, the two unique dataset have low mixing in (*c*) while the other cell types have high mixing. Although there is no labelled B cells, scANVI does not cluster B cells from the PBMC8K dataset with other celltypes in PBMC-CITE. The three metric we use to evaluate scANVI performance is minimally affected by the increase of number of classes.

**Figure 20:**
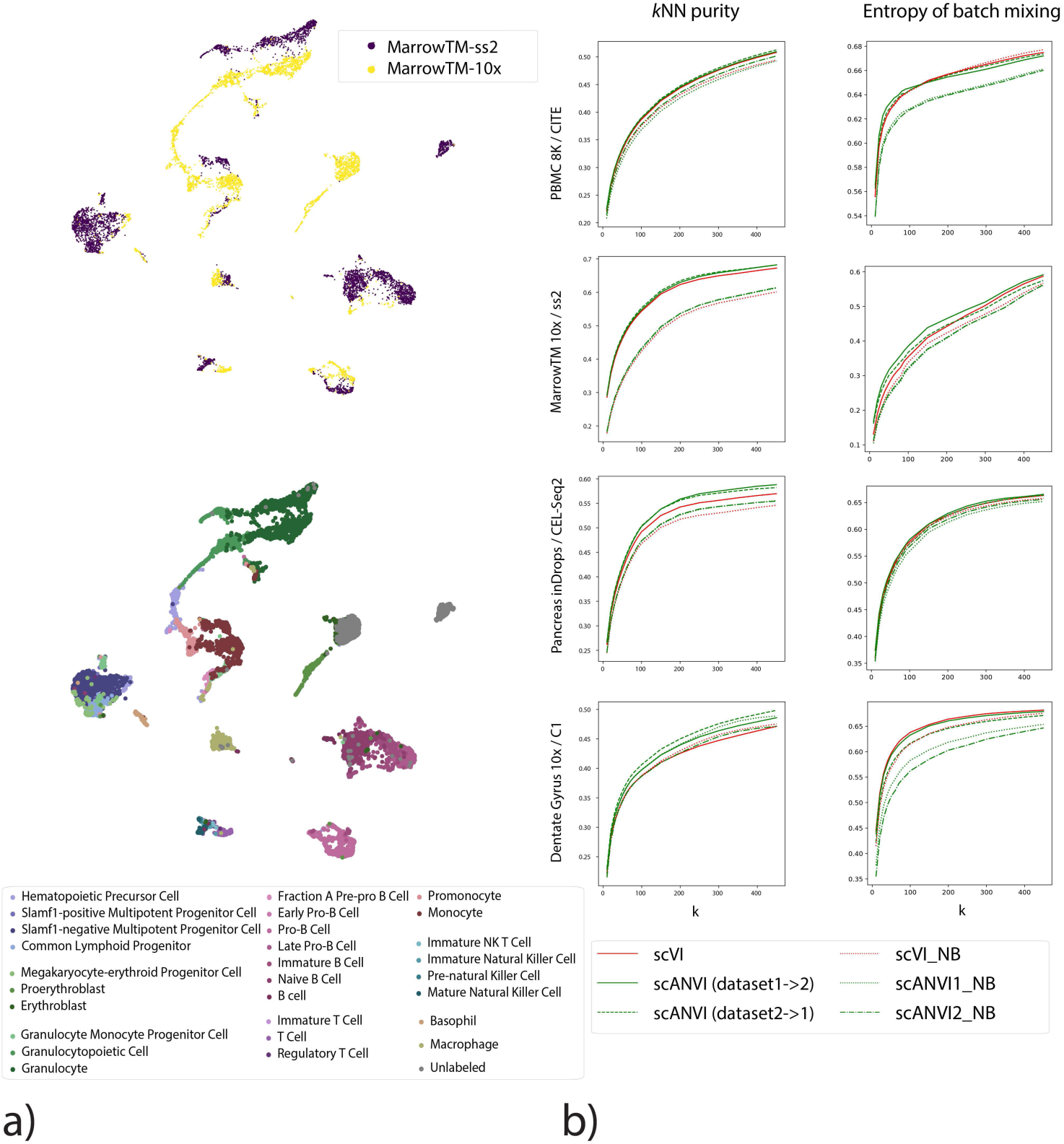
Performance of scVI and scANVI with a negative binomial (NB) distribution. (*a*) UMAP plot of the MarrowTM pair using a NB distribution for scVI. (*b*) Harmonization statistics and differences between regular scVI and NB version (scVI-NB). Dotted lines represent results using scVI-NB.

**Figure 21:**
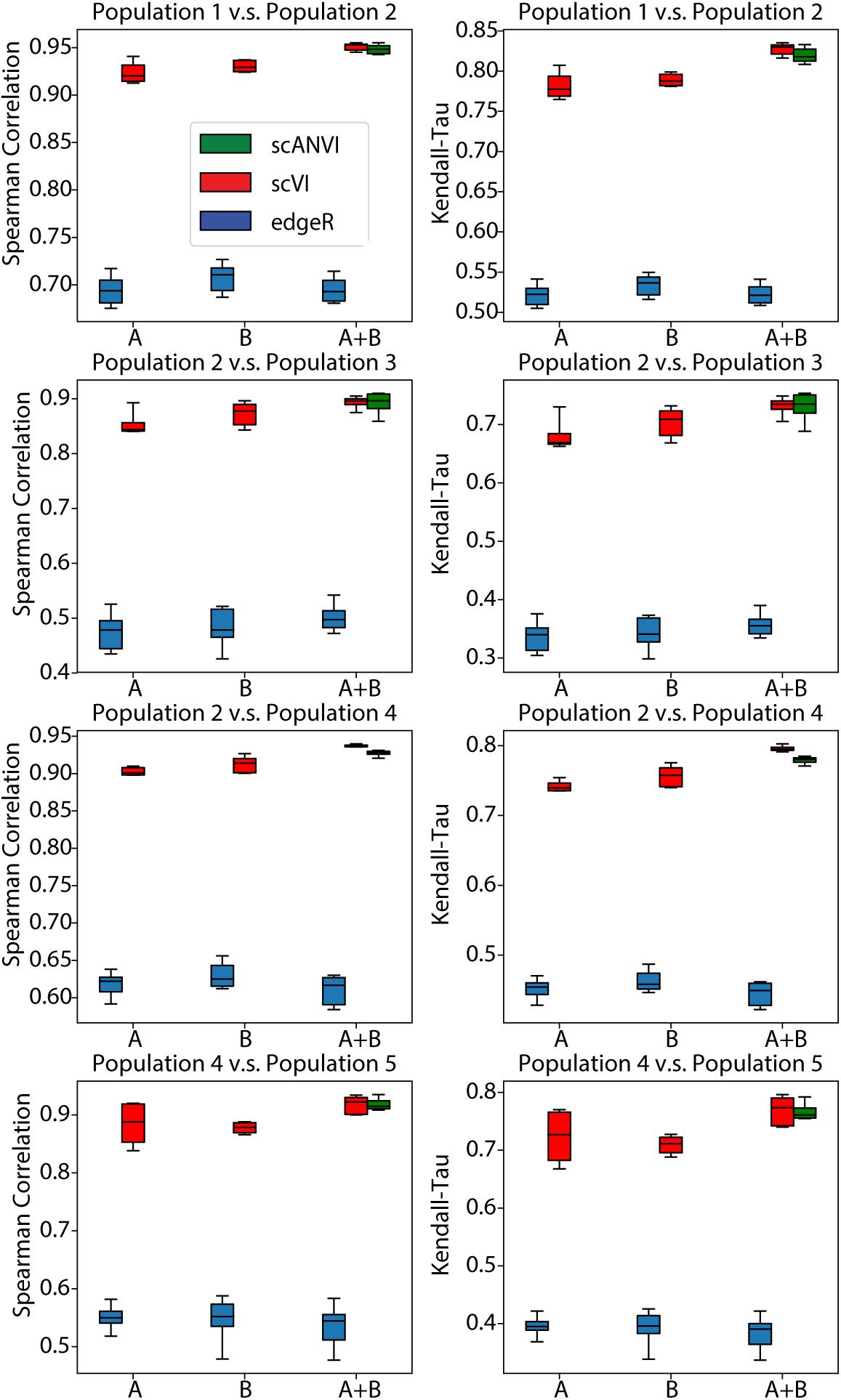
Differential Expression with a negative binomial version of scVI. We report all metrics on all pairs of cell types using the simulated dataset previously analyzed as in Supplementary Figure 16a.

## Supplementary Information

**Table 1:**
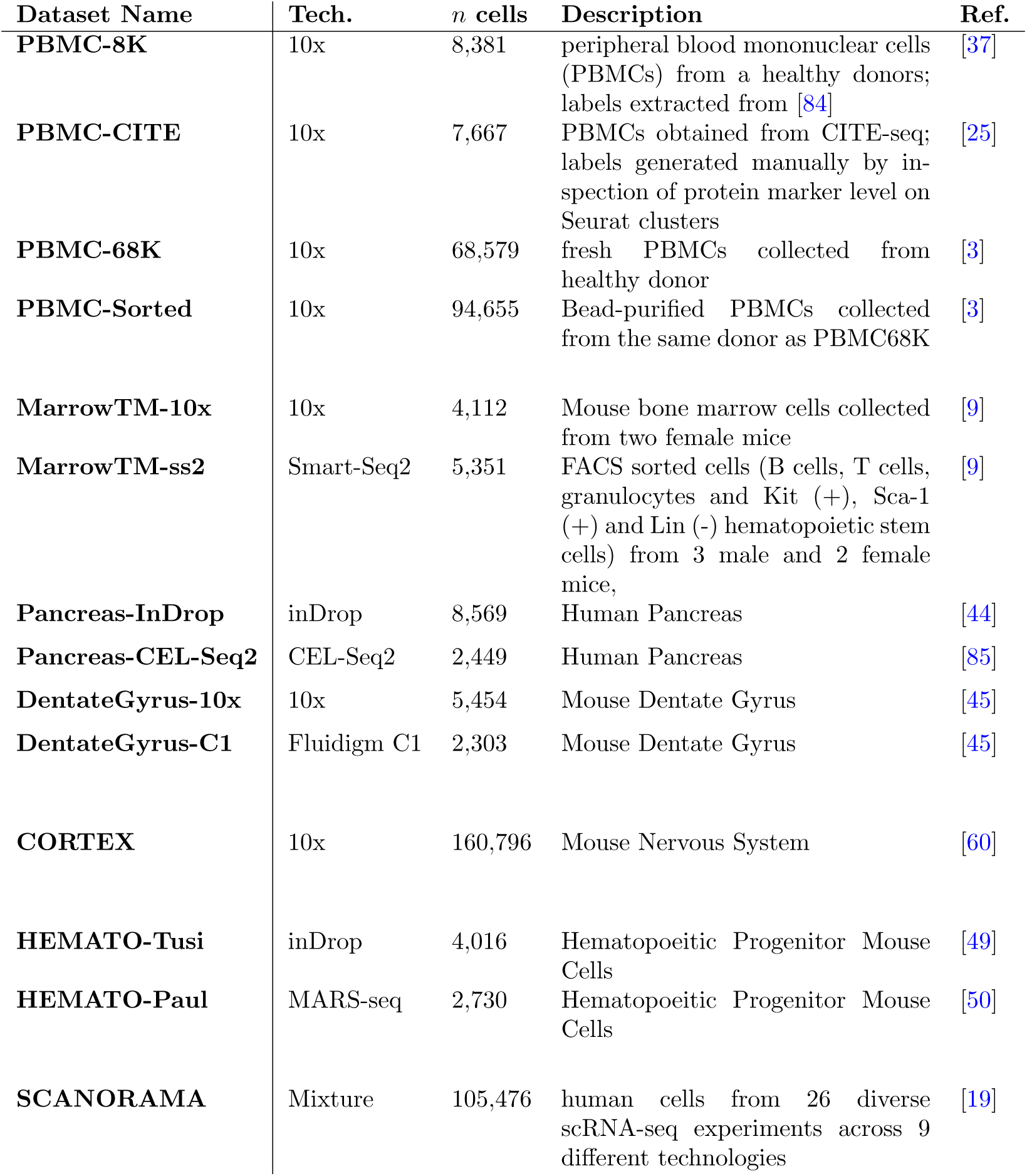
List of dataset used in this paper. Note that for the PBMC-Sorted 11 cell types were collected according to the paper but only 10 are available from the 10x website [37].

| Dataset Name | Tech. | *n* cells | Description | Ref. |
| --- | --- | --- | --- | --- |
| **PBMC-8K** | 10x | 8,381 | peripheral blood mononuclear cells (PBMCs) from a healthy donors; labels extracted from [84] | [37] |
| **PBMC-CITE** | 10x | 7,667 | PBMCs obtained from CITE-seq; labels generated manually by inspection of protein marker level on Seurat clusters | [25] |
| **PBMC-68K** | 10x | 68,579 | fresh PBMCs collected from healthy donor | [3] |
| **PBMC-Sorted** | 10x | 94,655 | Bead-purified PBMCs collected from the same donor as PBMC68K | [3] |
| **MarrowTM-10x** | 10x | 4,112 | Mouse bone marrow cells collected from two female mice | [9] |
| **MarrowTM-ss2** | Smart-Seq2 | 5,351 | FACS sorted cells (B cells, T cells, granulocytes and Kit (+), Sca-1 (+) and Lin (-) hematopoietic stem cells) from 3 male and 2 female mice, | [9] |
| **Pancreas-InDrop** | inDrop | 8,569 | Human Pancreas | [44] |
| **Pancreas-CEL-Seq2** | CEL-Seq2 | 2,449 | Human Pancreas | [85] |
| **DentateGyrus-10x** | 10x | 5,454 | Mouse Dentate Gyrus | [45] |
| **DentateGyrus-C1** | Fluidigm C1 | 2,303 | Mouse Dentate Gyrus | [45] |
| **CORTEX** | 10x | 160,796 | Mouse Nervous System | [60] |
| **HEMATO-Tusi** | inDrop | 4,016 | Hematopoietic Progenitor Mouse Cells | [49] |
| **HEMATO-Paul** | MARS-seq | 2,730 | Hematopoietic Progenitor Mouse Cells | [50] |
| **SCANORAMA** | Mixture | 105,476 | human cells from 26 diverse scRNA-seq experiments across 9 different technologies | [19] |

**Table 2:** Additional metric for retainment of structure via *k*-means. For scANVI we perform semi-supervision using the cell type label (not *k*-means cluster labels) from only one of the two datasets. Thus we train two separate models SCANVI1 and SCANVI2. To obtain a measure of clustering conservation, we first run *k*-means clustering in the latent space of dataset 1, then in the harmonized latent space using only cells from dataset 1. We compute the adjusted Rand Index of the two clustering results. We then do the same for dataset 2 and the final score is the average for both datasets.

| Method | PBMC8KCITE | MarrowTM | Pancreas | DentateGyrus |
| --- | --- | --- | --- | --- |
| scVI | 0.73395 | 0.74325 | 0.81425 | 0.5418 |
| scVI_nb | 0.7648 | 0.71035 | **0.8281** | **0.5915** |
| scANVI1 | 0.74465 | 0.7418 | 0.76625 | 0.55875 |
| scANVI2 | **0.8184** | **0.77825** | 0.78815 | 0.54135 |
| Seurat | 0.7351 | 0.72095 | 0.6174 | 0.4149 |
| MNN | 0.7364 | 0.6783 | 0.76205 | 0.4296 |
| PCA | 0.59895 | 0.6089 | 0.5474 | 0.4179 |

**Table 3:**
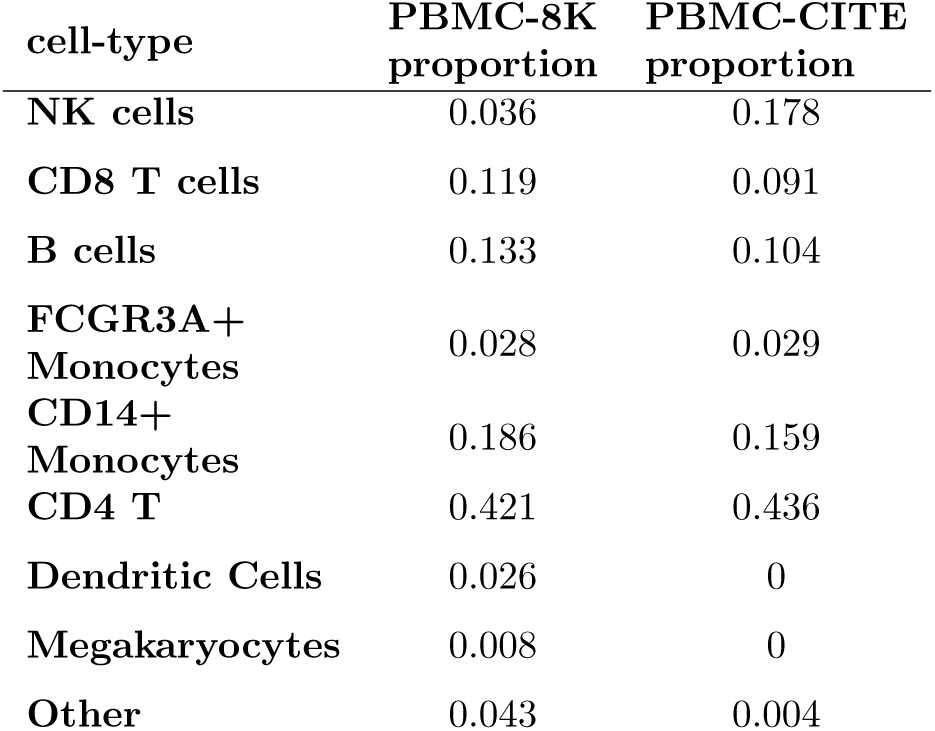
Compositions of cell-types in the PBMC-8K and the PBMC-CITE dataset

**Table 4:** Cell types present in the PBMC-sorted dataset.

| Cell Types | # cells |
| --- | --- |
| B cells | 10,085 |
| CD14+ Monocytes | 2,612 |
| CD34+ cells | 9,232 |
| CD4 T cells | 11,213 |
| CD56 NK cells | 8,385 |
| CD8 T cells | 10,209 |
| Memory T cells | 10,224 |
| Naive CD8 T cells | 11,953 |
| Naive T cells | 10,479 |
| Regulatory T cells | 10,263 |

## Supplementary Note 1 Related machine learning literature

Our approach relates to the machine learning literature in two major aspects. First, we will relate the problems of harmonization and annotation to the litterature of domain adaptation or covariate shift. Second, we will present all the different options for performing semi-supervised learning with variational autoencoders (VAEs) and further explain how they relate to scANVI.

### Domain adaptation

In the supervised learning framework, data is drawn from a distribution 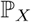 and one posit a conditional distribution 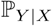 from which to draw the labels. Domain adaptation works in the setting where one might observe some paired data on a certain source domain 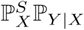 but no labels on a target domain 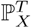 with 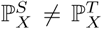 (note that some other flavors of domain adaptation problems make different hypothesis). This discrepancy between distributions can be referred to as a *covariate shift* in the statistical literature. Our problem of single-cell annotation is much related to this variant of supervised domain adaptation where one observe the cell labels for one dataset and which to transfer it to another (annotated) dataset. Let us note that this is a well established problem in the computer vision literature, where algorithms such as NBNN [36] and JDA [86] are used to transfer labels over datasets of images (e.g., with different lightning conditions, collections of images). As these algorithm might not scale to millions of samples, a notable contribution is the DANN method [35] which learns a adversarial classifier (justified by a theory of domain adaptation based on the **𝓗**-divergence). Another notable contribution is the variational fair autoencoder, which focuses on semi-supervised domain adaptation with VAEs [31] by adding a maximum mean discrepency loss between source and target latent distributions (see **Supplementary Note 2** for a discussion of adversarial or MMD-based methods). Moving away from the supervised domain adaptation scenario, recent work based on generative adversarial networks [87] focuses on unsupervised domain adaptation of *unpaired* datapoints, which is also the case in scRNA-seq where never observes cells from several batches. CycleGAN [46] is based on the idea of cycle consistency (a translator that would transform a french sentence into english and than back to french again should be the identity map). This idea was then improved by Cycada [88], which adds more consistency to the objective function. StarGAN [89] extended CycleGAN to multiple domains while reducing the overall complexity. Finally, MAGAN [47] proposed to add a correspondence loss to further orient the manifold alignment and facilitate the inference. Notably, MAGAN [47] was also applied to merging scRNA-seq and CyTOF data.

### Semi-supervised learning with variational autoencoders

Extending scVI to semi-supervised learning took some design that we describe here. Our first attempt was based on the M2 model [33], and is still implemented in our codebase^1^. While this algorithm performed a posteriori as good or better than than the final version of scANVI (in terms of cross-validation estimate of cell type prediction accuracy), it restrained our latent space *z* to be conditioned on the cell type label (as in the conditional VAE [90]) and it was not possible anymore to visualize the latent space, which is an crucial practice in scRNA-seq data analysis [91]. We therefore turned to extending scVI based on the M1 + M2 model [33], which enabled both a flexible modeling of cell types and visualization of a joint latent space for all cells. We also tried more complicated models based on the M1 + M2 model such as ADGM [34] and LVAE [92] but did not find them to contribute significantly enough to the accuracy, which might be either due to the labeling errors in the dataset, because dividing cells into cell types is a relatively easy problem in most regimes (e.g., not considering rare cell types) or because of the limited sample size in the datasets we consider.

## Supplementary Note 2 Statistical trade-offs in scRNA-seq harmonization

Harmonization is a hard and ill-defined problem. Especially, it can be difficult to formulate exactly its objective and at which level of granularity the “harmonization” is expected. Let us take the example of two PBMCs datasets that are exact biological replicates with same experimental conditions. In that case, the problem is well defined since we have a non-confounder assumption. Removing in a principled way all the variation in the data that corresponds to batches is reasonable. However, let’s take a slightly trickier case, one patient and one control dataset. If we look at the exact same metrics for harmonization, the resulting algorithm could discard any interesting biology that happens to the cells that get frozen. There is a confounder and the problem is much more difficult (ill-posed).

Therefore, it is extremely important to state how different models perform harmonization and what are their underlying hypothesis and modeling capabilities. For example, scmap [28] proposes a new gene filtering method to select gene that are claimed to be invariant to batch-effects. However, it is clear that over filtering can lead to ignoring biological information. MNNs [12] on its end assumes that the topology of cell types can be easily resolved by a *k*-nearest neighbor graph using a 𝕃^2^ normalization and distance scheme. This allows MNNs [12] and all the neighbor-matching-based approaches [20, 19] to remarkably merge batches with the risk of also merging cell types in the case where they are not detectable with this normalization scheme.

scVI and scANVI perform harmonization by learning a common generative model for a collection of gene expression probability distributions [*p*(*x* | *z, s*)]_*s*∈{1,…, *K*}_ indexed by the dataset-identifier *s*. The statistical richness of the collection of conditional variables [*p*(*x* | *z, s*)_*s*∈{1,…,*K*}_ dictates the flexibility of our model towards integrating datasets.

One notable factor that sensibly contributes to harmonization capabilities is the prior for cellspecific scaling factor *l*_*n*_ that is dataset-specific. This helps probabilistically removing library-size caused discrepancies in the measurements and is more principled than normalization of the raw data [93]. Another important parameter is the parametrization of the neural network that maps variables (*z, γ*) to the expected frequencies 𝔼[*w* | *z, s*] = *f*_*w*_(*z, s*). Since our function *f*_*w*_ is now potentially non-linear, our model can benefit from more flexibility and fit batch-specific effects locally for each cell types or phenotypical condition. Especially, depending on how one designs the neural architecture of *f*_*w*_, it is possible to more flexibly correct dataset-specific effects. More specifically, we treat *f*_*w*_ as a feed-forward neural network for which at each layer we concatenate the batch-identifier with the hidden activations. A consequence of this design is that with more hidden layers in *f*_*w*_, less parameters are shared and the family of distributions [*p*(*x* | *z, s*)]*s*∈{1,…,*K*} has more flexibility to fit batch-specific effects (Supplementary Figure 2a-c). Conversely, when the latent dimension grows, the generative network has less incentive to use the dataset covariate and might mildly duplicate the information in its latent space (Supplementary Figure 2d-f). Throughout the paper, we fix those parameters (**Online Methods** and show competitive performance for all our datasets.

Another insight comes from how the variational distribution *q*(*z* | *x, s*) is parametrized. Our neural networks plays the role of an explicit stochastic mapping from the gene expression *x*_*n*_ of a single-cell *n* to a location in a latent space *z*_*n*_ (a standard theme in scRNA-seq analysis, e.g LIGER [18], Seurat [13], scVI [29]). Via this map, we match an empirical distribution in gene expression space (i.e a dataset) 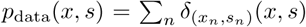, with a certain distribution on the latent space 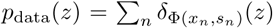. In certain cases, even though we designed this latent space to represent biological signal, the transformed dataset might still be confounded by technical effects [94]. In particular, this effect is susceptible to be severe when the generative model is not flexible enough to fit the dataset-specific effects or has a model misfit (as in, to some extent, the case with the CCA of [13] that assumes the data is log-normal). In this case, the go-to method is to empirically constrain the mapping (i.e the variational network in our case or the latent space itself in the case of SEURAT) to match the collections of variables *p*_data_(*z, s*)_*s*∈{1,…,*K*}_. However, each of these correction approaches has underlying modeling assumptions. For example, SAUCIE [21] and MMD ResNet [22] both propose to correct the variational distribution by adding on their objective function a non-parametric measure of distance between distributions (maximum mean discrepancy [95]). This approach specifically assumes that each dataset has the same cell type composition and the same biological signal and which is not reasonable in other cases than biological replicates. Seurat Alignment [13] on its end relies on a milder assumption that there is a common signal exactly reproducible between the two datasets and that CCA capture most of the biological variation, which is not obvious true considering limited suitability of linear Gaussian model for scRNA-seq. Seurat anchors [20] relies on CCA and suffer from the same problems as MNNs with its specific normalization scheme. Finally, the recent LIGER [18] method at first sight seems like a non-probabilistic version of scVI since it also learns a degenerate conditional distribution via its integrative non-negative matrix factorization [18] (NMF is a noisy-less version of a Gamma-Poisson generative model). However, it has a further quantile normalization of the latent spaces within clusters. First, the output of the clustering algorithm is not necessarily correct and could perturb downstream analyses. Second, if a cell type would be slightly different from one condition to another, this information would be lost in the final latent space. Overall, all these correction methods can therefore potentially lead to over-correction and statistical artifacts [32].

## Supplementary Note 3 Training Information

In order to train scanVI properly, several options are possible to train all the parameters (*θ* from the generative model, *φ* from the variational distribution and *θ*^𝓒^ from the labels’ posterior exclusively). In all cases, parameters *θ* and *φ* should minimize the evidence lower bound 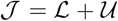 (decomposed over the labeled samples *L* and unlabeled samples 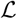). Furthermore, [33] suggests to optimize jointly a modified objective that penalize the ELBO by a classification loss *C* on the labeled data so that the parameters *θ*^C^ also benefits from the learned data. In [96], we interpreted this extension of the ELBO as a correction term for noisy labels. In this manuscript, we propose a specific training procedure for scANVI, based on alternating minimization on the parameters (*θ*, *C*) and *θ*^C^. A joint training, conform to the one suggested in [33] is described in **Algorithm 1**. Instead, an alternate training procedure would perform alternating minimization as described in **Algorithm 2**. The advantage of joint training is presumably that it shapes the latent space directly, through the modification of the encoder’s weights. The advantage of the alternate training, is that the learning of the cell type edge converges properly at the end of each epoch, and improve indirectly the latent space quality, through the next steps of optimization. We have noticed in practice that using the joint training in the case of transferring labels from one dataset to another might break down the mixing in the latent space, as the loss *C* has only contributions from a unique dataset. Therefore, throughout the paper, we consistently used the alternating procedure of **Algorithm 2**.

### Algorithm 1: Train jointly

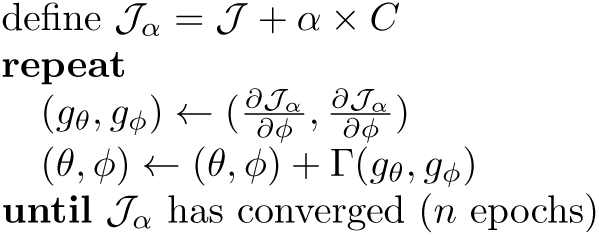

### Algorithm 2: Train alternately (Default scANVI)

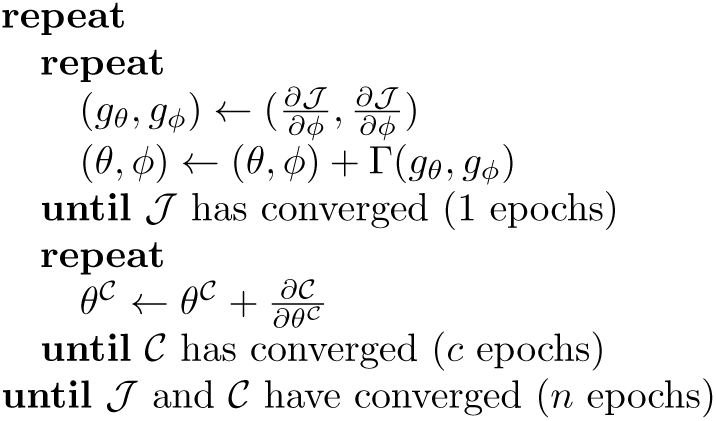

## Supplementary Note 4 Alternative model choices

### 4.1 Zero-inflation in the context of harmonization

In this manuscript, we mainly rely on a zero-inflated negative binomial (ZINB) distribution for the counts — which is a widely used model for single cell transcriptomics data. Recent research however suggest that for some technologies such as droplet-based UMI single-cell protocols, the abundance of zero mainly may be explained only by limited sensitivity and subsampling effect [97]. On the other hand, zero-inflated might be a realistic addition to count distributions for describing full-length plate-based technologies data [98]. Indeed, it has been hypothesized that PCR duplication or uneven fragment sampling may be responsible for zero-inflation in plate-based technologies [97]. Still, no statistically significant conclusion can be drawn about whether negative binomial distribution better fits UMI technologies while ZINB is needed for full-length method. Although full-length methods seem more affected by technical noise, full-length sequencing provides extra information at the transcript resolution that is lost in droplet-based UMI methods. This is why many consortia have chosen to obtain data by both full-length and UMI methods [9, 99], and why a method that can merge these two data types is crucial for analyzing single cell transcriptomics data. scVI and scANVI are both flexible to use both ZINB and NB methods and we show in Supplementary Figure 20 that the benchmarking results are similar using both methods in all four datasets used in our benchmarking procedure. However, we observed that for MarrowTM dataset, using the ZINB version of scVI does perform better in terms of harmonization. Since in this case, we are merging a 10x dataset with a Smart-Seq2 dataset, we compared the average zero-inflation parameter for each gene in each dataset and found that the differences between Smart-Seq2 and 10x are significantly skewed to the right (more zero-inflation in Smart-Seq2, *p*=6.3e-31, using a D’Agostino-Pearson *K*^2^ test). Finally, we also performed differential expression with negative binomial versions of scVI and scANVI and show that results are similar (Supplementary Figure 21). These results show that both NB and ZINB model can be used in scVI and scANVI and in most instances, downstream analysis might not be impacted by this modeling choice. Interestingly, our model successfully learns the difference in the degree of zero-inflation in different datasets while merging them and exploits this information when necessary.

### 4.2 Library-size prior for scVI and scANVI

Besides zero-inflation, another major difference between different sequencing technologies is the library size. scVI and scANVI infers library size from the data with a batch-specific prior. To evaluate the effectiveness of both the model and the prior choice, we compute the negative log likelihood of the scVI model using ZINB with batch-specific library size prior, ZINB with shared library size prior, NB with a batch-specific library size prior, and NB with shared library size prior. All four models are fit on two pairs of datasets, DentateGyrus (Fluidigm C1 and 10x) and MarrowTM (10x and SmartSeq2). Since only the second pair is a comparison between UMI and full-length datasets, we expect the differences between the four model choices to be larger in the MarrowTM comparison, and that ZINB with batch-specific library size prior to have the highest likelihood (reported in the table below).

**Table:** Assessing the model’s fit with different configuration on two pairs of dataset

| Negative log-likelihood | DentateGyrus | MarrowTM |
| --- | --- | --- |
| **ZINB model / batch-specific prior** | 514.5 | 2339.8 |
| **ZINB model / shared prior** | 513.8 | 2349.2 |
| **NB model / batch-specific prior** | 523.4 | 2531.3 |
| **NB model / shared prior** | 524.0 | 2544.1 |

## Supplementary Note 5 Hierarchical classification

In this note, we explain the modeling details for an extension of scANVI where the cell types annotation can follow a two-level hierarchical structure. Let us note that this can in principle be adapted to any arbitrary tree representation of cell types taxonomy. This taxonomy needs to be hard-coded and known *a priori*.

We do not need to formally modify the graphical model but only how we structure the variable *c*_*n*_. Notably, we formally pose 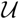 and consider the original Bayesian network as an hypergraph. In the generative model, the conditional distribution *p*(*z* | *u, c*) = *p*(*z* | *u, y*). Since the cell type group indicator variable *y*^*g*^ is not as refined than the cell type itself *y*.

However, what sensibly changes is the parametrization of the variational distribution *q*(*c* | *z*) = *q*(*y, y*^*g*^ | *z*). The prior taxonomy knowledge encapsulates whether the assignment (*y*^*g*^, *y*) = (*i, j*) is biologically possible (i.e cell type *i* is a sub-population of group cell type *j*). We encode this biological compatibility into a parent function *π*: {0, …, *K*} → {0, …, *K*^*g*^} that maps a cell type to its parent in the hierarchy. We note for simplicity 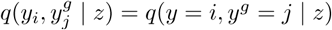. We then use two neural networks *f* and *f*_*g*_ (with softmax non-linearities) that map the latent space to the joint approximate posterior *q*(*y*, *y*^*g*^ | *z*) with the following rules:

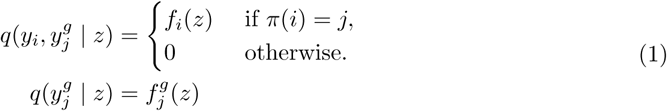

Then, we can derive the marginal probability over finer cell types classes using the chain rule and Bayes rule:

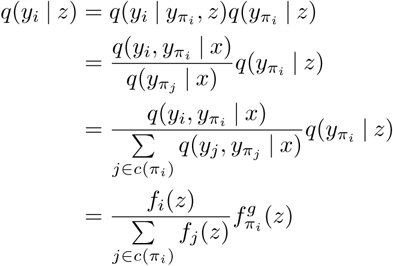

where *c*(*π*_*i*_) denotes the set of children of node *i* children.

## Supplementary Note 6 Evidence Lower Bound decomposition

For notational convenience, we do not write here that all the probabilities are conditioned on the batch identifier *s*. We also report the evidence lower bound (ELBO) only for one sample (i.e one cell) and drop the index notations by substituting {*x*_*n*_, *z*_*n*_, *u*_*n*_, *c*_*n*_, *l*_*n*_} by {*x, z, u, c, l*}. This is without loss of generality since all the cells are independent and identically distributed under our model. Bayesian inference aims at maximizing the marginal likelihood of our datapoints. We also drop the parameters Θ (resp. Φ) of the generative model (resp. the variational distribution) for notational convenience. We will derive the ELBO in the case where *c* is not observed (almost same calculations resolve the case where *c* is observed). Notably, we derive the ELBO using Jensen inequality weighted by the variational distribution *q*(*z, u, l, c* | *x*). Similar derivations can be found in the variational autoencoder literature (eg. [33]). We assume our variational distribution factorizes as:

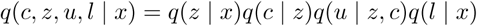

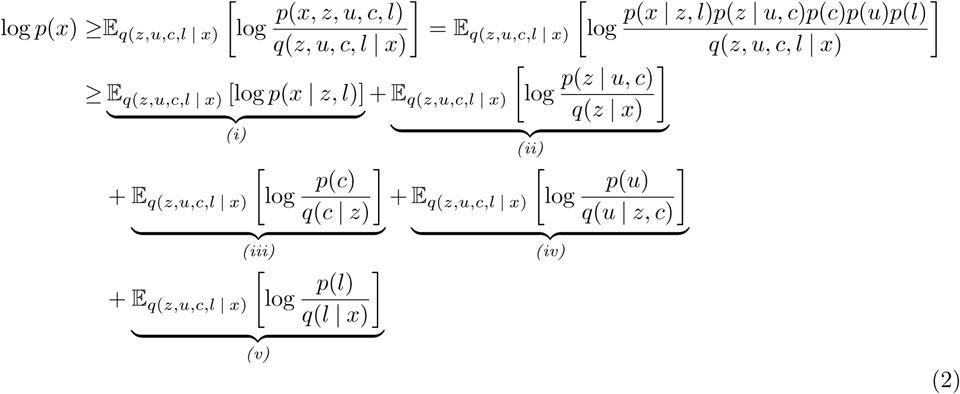

Then we simplify each individual term of the ELBO in 2 by recognizing KL divergence terms. In particular, we use subscript notation KL_G_ (*.||.*), and KL_M_ (*.||.*) to denote Gaussian and multinomial KL divergences.

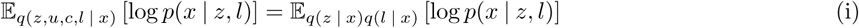

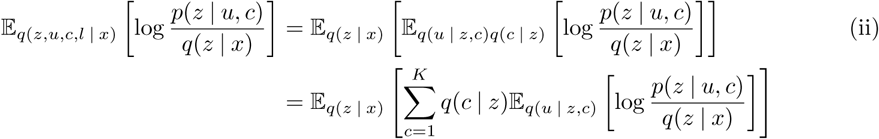

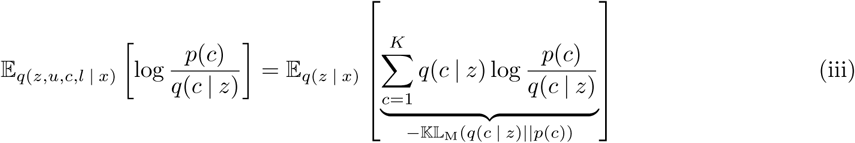

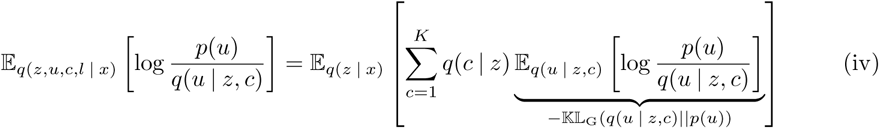

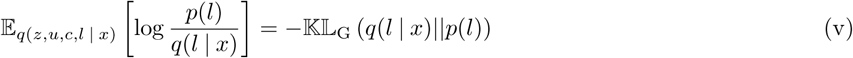

## Footnotes

1 VAEC: https://github.com/YosefLab/scVI/blob/master/scvi/models/vaec.py

